# Sequential, Chromosome-Specific Glutamine Synthetase Double Knockout with Cas-CLOVER Establishes Enhanced CHO Platforms for Cell Line Development

**DOI:** 10.64898/2026.01.06.698029

**Authors:** Cintia Gomez Limia, Hsueh Che Cheng, Daniel Machado, Thomas Hart, Mark Cameron McHargue, Chia-Chuan Cho, Corey Brizzee, Jack Crawford

## Abstract

Cas-CLOVER is an emerging high-fidelity genome editing system that enables precise and efficient cell engineering. In this study, we applied Cas-CLOVER to establish a robust, gene-edited platform in suspension-adapted CHO-K1 cells supporting cell line development (CLD) for biopharmaceutical production. An attractive strategy for high yield clone selection is the use of glutamine synthetase (GS) knockout CHO cells. The primary GS gene resides on chromosome 5 (GS5), while a recently identified GS pseudogene is located on chromosome 1 (GS1). To compare editing efficiency, we evaluated Cas-CLOVER and Cas9 at both GS loci using the Neon™ Transfection System. Cas-CLOVER achieved 84% editing at GS5 and 74% at GS1, markedly higher than Cas 9. Leveraging Cas-CLOVER’s dual-guide RNA design, we generated a GS5 single knockout (GS5-SKO) and subsequently a double knockout (GS-DKO) line at both the GS5 and GS1 loci, both with none detected off-target mutations analyzed in 40 predictably off-target sites. For functional validation, these cell lines were engineered with the proprietary Harbor-IN transposase system to stably express trastuzumab. Using an optimized protocol, the resulting GS-DKO platform, termed CleanCut GS CHO, enabled stringent selection and yielded high-producing clones with cell-specific productivity exceeding 100 pg/cell/day and antibody titers greater than 5 g/L in 24 deep well-plate fed-batch cultures after 14 days. The antibody titer stability analysis showed consistency over 60 generations. Collectively, these findings establish Cas-CLOVER as a versatile genome editing tool for developing high-yield CHO host platforms in CLD.

## 1. Introduction

Chinese hamster ovary (CHO) cells are the dominant expression system for recombinant proteins, particularly therapeutic antibodies, owing to their robust growth in suspension, adaptability to chemically defined media, capacity for human-compatible glycosylation, and high yield in fed-batch cultures ^1,2^. CHO cells serve as the cornerstone of cell line development (CLD) platforms for the production of life-saving therapeutics such as pembrolizumab (Keytruda®) ^3^, adalimumab (Humira®) ^4^, rituximab ^5^ and trastuzumab ^6^.

Efficient cell line development depends on the ability to identify high-producing CHO clones, yet this process is often prolonged by the need to isolate and evaluate large numbers of candidates. Strategies that enrich high-yield clones early in development streamline CLD workflows and accelerate the path to manufacturing. Two principal selection systems dominate CHO cell line development: the dihydrofolate reductase (DHFR) system ^7^, which relies on gene amplification under selective pressure, and the glutamine synthetase (GS) system, which leverages metabolic selection in glutamine-deficient conditions. In this study, we focus on the GS system due to its widespread industrial adoption and proven ability to generate high-yield clones. GS-based selection is typically implemented using methionine sulfoximine (MSX), an inhibitor of GS, in glutamine-free media. MSX enriches the cell population for high GS-expressing cells carrying an integrated GS transgene. While this selection method works in standard CHO cell lines, targeted knockout of the endogenous GS gene has been shown to enhance selection stringency ^8,9^.

One of the foundational studies on generating CHO cells with an endogenous GS gene KO on chromosome 5 (GS5) employed Zinc Finger Nuclease (ZFN) technology. The researchers confirmed successful GS disruption by demonstrating glutamine-dependent growth in all GS-KO cell lines. The authors showed that clones with biallelic GS5 disruption and loss of GS protein expression analyzed by WB were entirely dependent on exogenous glutamine for survival, consistent with a loss of GS enzyme activity ^8^. The selection system was enhanced by employing a weakened SV40E promoter to modulate GS expression in CHO cells ^10^. However, a low expression of pseudo-GS gene on chromosome 1 (GS1) and GS activity was reported ^11,12^.

As alternative, to generate CHO cell lines for selection capabilities and other CLD applications, CRISPR-based gene editing strategies have been implemented. CRISPR-based gene editing strategies were used to disrupt GS1 gene in addition to the major GS5. By simultaneously targeting both loci with CRISPR/Cpf1, double knockouts, named in the study as dGS5,1KO, were generated in CHO-S and CHO-K1. Notably, the CHO-K1 dGS5,1KO exhibited significant gains in titer and cell-specific productivity, a particularly important result given CHO-K1’s widespread use in stable GMP manufacturing, whereas CHO-S is more commonly employed for transient expression. However, these studies did not incorporate advanced stable gene delivery systems to increase productivity ^11^.

The rationale for generating GS double KO cells is based on the presence of both these GS genes expressed in CHO cells, with GS5 showing high expression and GS1 showing low expression. The highly expressed GS5 gene contains exons 6 and 7, which harbor the essential sequence required for GS enzymatic activity ^11^.In contrast, GS1 is weakly expressed and consists of only a single exon, and its sequence and expression have previously been confirmed through PCR amplification of its mRNA ^12^. GS1 sequence does not represent the complete GS coding region found on GS5, however, it retains the substrate-binding beta-grasp domain associated with glutamate-ammonia-ligase activity in the enzyme, indicating that some residual glutamine-synthetic function may persist. Therefore, in the current study we KO both GS5 and GS1 loci in CHO-K1 comparing the antibody production after glutamine selection in single vs double KO CHO cells pool, using Cas-CLOVER technology.

More recently, CRISPR/Cas9 has been applied to optimize CHO hosts through targeted genome editing. For example, knocking out Fut8 gene enables the production of afucosylated antibodies with enhanced antibody-dependent cellular cytotoxicity (ADCC) ^13,14^. Similarly, multiplex disruption of lactate dehydrogenases and pyruvate dehydrogenase kinases prevent lactate accumulation, thereby preserving growth and improving cell densities in fed-batch cultures ^15^.

Despite the widespread use of CRISPR/Cas9 for genome engineering, Cas9 lacks precision and has been associated with off target editing rates exceeding 10% in CHO cells ^16^. Undesired chromosomal rearrangements can also occur at frequencies of ∼7%, which is a particular concern for CLD as it complicates clonal selection and the transition from discovery to production ^17^. In addition, the complex and fragmented intellectual property landscape surrounding Cas9 has made many industrial biomanufacturers cautious to adopt it broadly, slowing the wider deployment of streamlined genome editing and limiting access to an otherwise enabling tool. An attractive next-generation alternative is Cas-CLOVER, a dimeric RNA-guided endonuclease that offers high-fidelity editing with markedly reduced off-target activity and has demonstrated successful application in mammalian systems. Madison et al. showed that indel frequencies at candidate off-target sites were undetectable or below 0.1% by GUIDE-seq and amplicon-seq analysis. Compared to CRISPR-Cas9, Cas-CLOVER exhibited 5- to 25-fold lower off-target rates and produced low levels of balanced translocations (<0.2%) with nearly undetectable unbalanced translocations as measured by AMP-seq ^18^.

The two main approaches for protein expressions in CHO cells are transient gene expression and stable gene expression. Several research groups showed stable expression can increase antibody production compared to transient expression thus supporting the use of transposase/transposon systems ^19,20,21^. For stable expression, transposase-based platforms, such as the Super piggyBac transposon system, represent some of the most efficient tools for stable CHO CLD, yielding multifold increases in antibody titers and specific productivity compared with random integration approaches ^21^. Transposase systems such as piggyBac also enable the delivery of large genetic cargos and preferential integration into transcriptionally active open chromatin sites, thereby improving genomic safety and ensuring stability across generations. Here we introduce Harbor-IN, a proprietary transposase expressed as codon-optimized mRNA and manufactured using *in vitro* transcription (IVT) methods to enhance stability and efficiency. Unlike other transposase systems, Harbor-IN is not licensed restricted and is commercially available as a reagent, offering biomanufacturers a practical and accessible solution for stable CHO cell line development.

To our knowledge, the Cas-CLOVER dimeric nuclease system has not previously been applied to sequentially target two highly homologous genes in CHO cells, and prior reports of the dGS5,1KO cell line did not incorporate advanced CLD workflows using transposase systems to enhance titer and productivity^11^. The objective of this study was therefore to assess the efficiency and specificity of Cas-CLOVER in generating a GS-double knockout (GS-DKO) in CHO-K1, termed as CleanCut GS, and to stably integrate therapeutically relevant genes using a proprietary Harbor-IN transposase. While our CLD processes have traditionally relied on the 4D Nucleofector with consistent success, optimization of Cas-CLOVER editing on the Neon Transfection system achieved >90% editing efficiency, underscoring both the versatility and robustness of Cas-CLOVER. We further demonstrate that CleanCut GS CHO cells, when engineered with Harbor-IN to express trastuzumab, supported stable antibody production with titers exceeding 5 g/L and cell-specific productivity (Qp) above 100 pg/cell/day, with consistent stability of titers over 60 generations. Collectively, these studies demonstrate that Cas-CLOVER enabled the generation of CleanCut GS CHO, a versatile and stable platform that can be deployed either individually or as an integrated package with Harbor-IN transposase to enable cost-effective, next-generation biologics manufacturing. CleanCut GS CHO pools expressing trastuzumab deliver stable, high-titer production on an accelerated timeline, lowering manufacturing costs while rapidly supplying material for development. Compared with immediately generating single-cell clones (SCC), stable CHO bulk pools expressing the antibody of interest offer faster production, screening, and expansion effort. Furthermore, stable CHO bulk pools enable greater process representativeness by averaging across many producers and incorporate sufficient stability to streamline early media and feed optimization which allows for straightforward scalability to pilot and manufacturing scales.

## 2. Material and Methods

### 2.1. Cas-CLOVER construct and gRNA design

Cas-CLOVER was originally engineered by fusing the C-terminal nuclease domain of Clo051, a type IIS restriction endonuclease from the bacteria *Clostridium* to the N-terminus of catalytically inactive Cas9 (dCas9) via a flexible GGGGS (G4S) linker. Demeetra’s Cas-CLOVER mRNA is manufactured using a proprietary process which enhances stability and activity (Demeetra, Cat# R08). Paired guide RNAs targeting exon 6 of the GS gene on GS5 and exon 1 on GS1 (homology sequence in the junction exon 4/5 with GS5) were designed and synthesized with 20 bp recognition sequences and 17 or 16 bp spacers between each guide pair, respectively (Table 1). The design was validated through bioinformatic pipelines to minimize off-target activity by CrisPick from Broad Institute software.

**Table 1.**
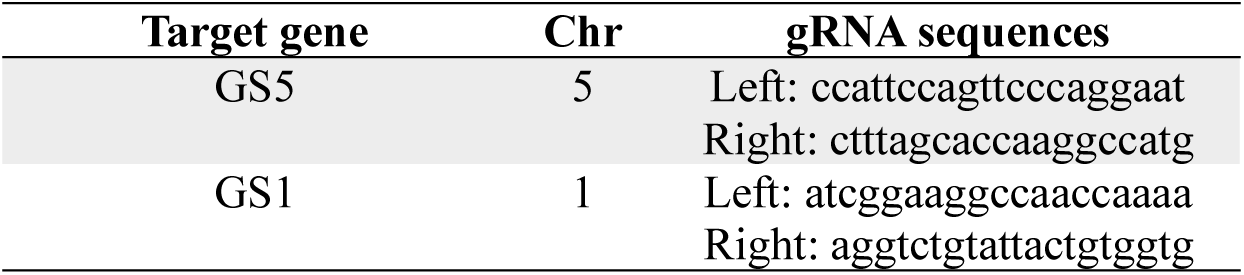
Summarization of gRNAs gene editing used to flank the target region and disrupt the GS5 or GS1 genes.

### 2.2. Harbor-In Plasmid construction and mRNA reagents

An empty transposon plasmid (Demeetra, Cat# D01) containing a multiple cloning site (MCS), proprietary core insulators, and Harbor-IN inverted terminal repeats (ITRs) were used as the cloning backbone. Trastuzumab expression was driven by separate transcriptional units: the light chain (LC) under control of a cytomegalovirus (CMV) promoter, and the heavy chain (HC) under an enhancer/promoter complex consisting of the CMV enhancer, Ef1α core promoter, and a 5′ HTLV element. For GS selection, the GS transgene was placed under a weakened SV40E promoter (Figure S1). Harbor-IN mRNA (Demeetra, Cat# R07) was generated by *in vitro* transcription (IVT) from a plasmid template containing a T7 promoter and the optimized Harbor-IN coding sequence.

### 2.3. Cell Culture

Adherent CHO-K1 cells were thawed and maintained using standard conditions. Cells gradually underwent adaptation from adherent culture to suspension in chemically defined CD FortiCHO^TM^ medium (Gibco, Cat#A1148301) containing 8 mM L-glutamine (Gibco, Cat#25030-081). Adaptation was initiated in T-75 flasks with 0.05% trypsin-EDTA (Gibco, Cat#25300-062) and performed sequentially by stepwise dilution of Ham’s F-12 medium (Sigma-Aldrich, Cat#N4888) with increasing proportions of CD FortiCHO^TM^ medium (25:75, 50:50, 75:25, 90:10, 95:5, 97.5:2.5), followed by 100% of CD FortiCHO^TM^ medium. Suspension-adapted CHO-K1 cells were cultured in fresh CD FortiCHO^TM^ medium in complete absence of L-glutamine and aggregative score 0 (no clumps). After successful adaptation, suspension CHO-K1 cells were maintained in fresh CD FortiCHO™ medium in the complete absence of L-glutamine. Cultures were grown in unbaffled 125-mL Erlenmeyer (E-125) shake flasks with a 30 mL working volume and incubated at 37 °C with 8% CO₂ on an orbital shaker at 125 rpm. Suspension cultures were passaged every 4 days by inoculating into fresh medium at 0.3 × 10⁶ viable cells/mL; cell density and viability were confirmed prior to each passage with an EVE^TM^ automatic cell counter (NanoEntek).

### 2.4. Cas-CLOVER, CRISPR/Cas9 and Harbor-IN Transfection

To test editing efficiencies at the GS5 and GS1 loci, suspension CHO-K1 cells were transfected using the Neon^TM^ NxT Electroporation System (Thermo Fisher Scientific, NEON1S). Cells in exponential growth with viability >90% were harvested, washed in Ca²⁺/Mg²⁺-free PBS, and resuspended in Buffer R. For each condition2.5 × 10⁵ cells were electroporated with Cas-CLOVER or Cas9 mRNA and gRNAs at a 1:2 ratio using 10 µL Neon tips at 1,650 V, 10 ms, and 3 pulses. Immediately after electroporation, cells were transferred into pre-warmed, antibiotic-free CD FortiCHO^TM^ medium and placed in incubator at 37 °C, 8% CO₂. Viability was assessed using the Trypan Blue Stain 0.4% dye exclusion assay to identify and count viable and non-viable cells (Thermo Fisher Scientific, Cat#15-250-061). Cells were seeded in 24-well low-attachment plates immediately after transfection and incubated for 72 h; cell pellets were then harvested for analysis by T7E mismatch-detection assay. The experiment was performed in biological triplicates.

CleanCut GS CHO cells were generated by transfecting the parental cells using Lonza 4D-Nucleofector System. 2 × 10⁶ cells were resuspended in 100 µL of SF buffer - Cell Line 4D-Nucleofector X Kit (Lonza, Cat# V4XC-2032) with 2.5 µg of Cas-CLOVER mRNA and 2.5 µg of each gRNA or 2 µg of Harbor-IN transposase and 8 µg of transposon plasmid, following Program DS-137. Immediately following nucleofection, cells were transferred from cuvette into pre-warmed CD FortiCHO^TM^ medium supplemented with 8 mM L-glutamine and placed into incubator at 37 °C, with 8% CO₂.

### 2.5. T7 Endonuclease I (T7EI) assay

Genome editing efficiency was assessed using the EnGen^®^ Mutation Detection Kit (NEB, Cat# E3321S). After DNA extraction, 100 ng of genomic DNA was used to perform target specific PCR, the corresponding amplicon was quantified by Nanodrop. Approximately 200 ng of purified product was denatured at 95 °C and gradually cooled to form heteroduplexes following the instruction with small modifications. Samples were digested with T7EI at 37 °C for 15 min, followed by Proteinase K treatment at 37°C for 5 min. The samples were run on a Novex™ TBE Gels, 4–20% (Thermo Fisher Scientific, Cat#EC6225BOX). Editing efficiency was determined by densitometry using the formula: % modification = 100 × [1 – (1 – fraction cleaved)^1/2^]. The primers used for the T7 assay are listed in Table 2.

**Table 2.**
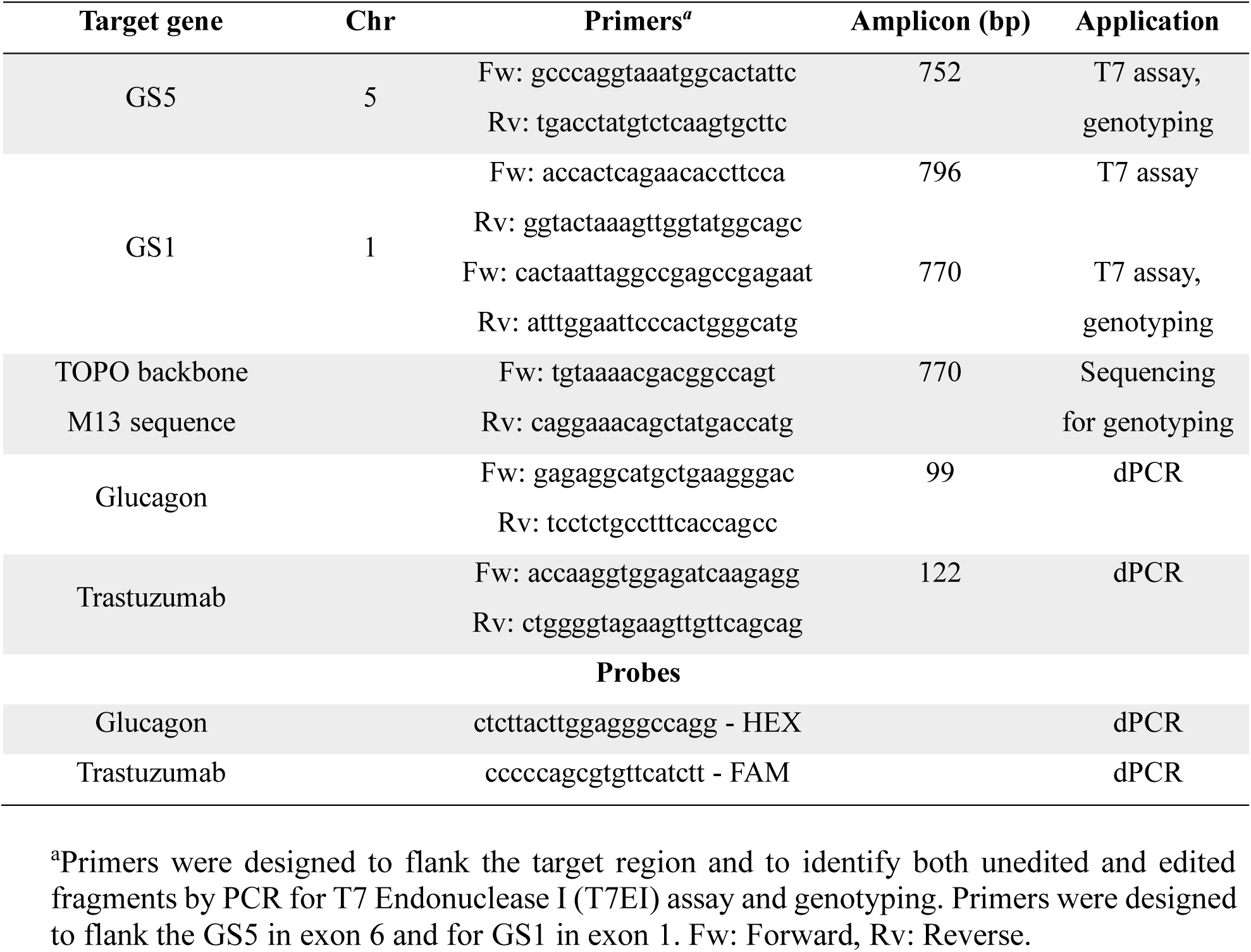
Primers and probes used for gene editing detection, genotyping validation or dPCR assay.

### 2.6. PCR for T7EI assay detection and genotyping validation

Genomic DNA was extracted from CHO cells pellet using the DNeasy Blood & Tissue Kit (Qiagen, Cat#69504). PCR amplification was performed with Q5® High-Fidelity 2X Master Mix (NEB, Cat#M0492S) following the manufacturer’s instructions. The amplicons were used for T7EI assay detection or for genotyping validation. The amplicons were cloned into TOPO^TM^ TA Cloning^TM^ Kit for sequencing (Thermo Fisher Scientific, Cat# K450030) following the manufacturer’s instructions. The colony screening was performed using the M13 primers with GoTaq® G2 DNA Polymerase (Promega, Cat#M7823), carrying out the reaction on Applied Biosystems 2720 (Thermo Cycler Model# 4359659). The amplicons were sequenced by Sanger sequencing with the M13 Fw primer. The primers used for genotyping are listed in Table 2.

### 2.7. Growth, viability assessment, and GS selection

Cells were expanded in Erlenmeyer shake flasks and evaluated for growth parameters including post-thaw viability ≥90%, recovery to ≥90% viability within ≤3 days, and doubling time (DT) ≤24 h using the following the formula: DT = (t2−t1)* (ln(2)/ ln(N1/N2)), N1, N2: viable cell density (VCD) at times t1 and t2 (days). Cell viability was assessed in CD FortiCHO^TM^ medium supplemented with or without L-glutamine to determine dependence on exogenous L-glutamine.

For stable GS selection, two days post-nucleofection cells were transferred at 2 × 10⁶ cells into T-25 low-attachment flasks containing 5 mL L-glutamine-free CD FortiCHO^TM^ medium and cultured at 90 rpm, at 37°C and 8% CO₂. On day 5–6, when cultures reached around 9 × 10⁶ cells with viability >90%, cells were scaled into 30 mL unbaffled E125 flasks at 0.3 × 10⁶ cells/mL. Selection was maintained for 14 days in L-glutamine-free medium, with reseeding and medium exchange at 0.3 × 10⁶ cells/mL every 2 or 3 days.

### 2.8. Single-Cell Cloning (SCC)

Sequencing-confirmed edited pools were subjected to SCC using the Verified In-Situ Plate Seeding (VIPS™) system (Solentim/Advanced Instruments). Cells were plated at one cell/well into 96-well plates containing 200 uL CD FortiCHO™ medium supplemented with 8 mM L-glutamine, and 5% InstiGRO™ CHO supplement (Advanced Instruments, Cat# RS-1125). Colony growth was monitored weekly for two weeks to assess plating efficiency and clonal outgrowth. Colony outgrowth was monitored by automated imaging to assess plating efficiency, monoclonality, and proliferation.

### 2.9. 24 deep well plate (DWP) fed batch for pool or clone and sample collection

For fed-batch studies, on day 0, cells were seeded at a density of 1 × 10⁶ cells/mL in a total volume of 3.8 mL with L-glutamine-free CD FortiCHO^TM^ medium in 24 DWP. PBS was added in all perimeter wells around the wells containing cells to avoid inconsistencies with cell aeration and shear stress. To maintain optimal conditions for cells growing an orbital shaker was used at 275 rpm. Beginning on day 2, cells were fed daily with a 2% (v/v) HyClone CellBoost™ mixture 7a and 7b supplement, (7a:7b) 10:1 (Cytiva, 7a, Cat# SH31119.02, 7b, Cat# SH31120.03) cells were fed every day from day 2 to day 13. On day 4, the temperature was shifted down from 37 to 32 °C. The 24 DWP was placed in a Deutz Clamp (EnzyScreen CR1702) and used a sandwich cover (EnzyScreen CR1224) to control evaporation rate and humidity. VCD and viability were measured by dye exclusion stained using Trypan blue. Supernatant was collected after centrifugation at 125 rpm for 7 min and stored at −80°C to measure antibody titer. Samples were collected on day 7, 9, 11, or 12, 13 and 14 for the experiments.

### 2.10. Titer assessment and specific productivity calculation

The supernatant collected on day 2, 7, 9, 11 or/and 12, 13 and 14 was diluted with medium 30- or 60-fold for titer quantification. IgG concentrations in culture supernatants were quantified using the ValitaTiter assay (Beckman Coulter, Cat. VAL003) according to the manufacturer’s instructions. A calibration curve was prepared from the kit standards covering 2.5 to 100 mg/L. Standards, samples, and kit-provided quality control were dispensed into a black 96-well plate in triplicate. After addition of the ValitaTiter reagents, plates were mixed gently and incubated for the recommended time at room temperature. Fluorescence polarization was read on a compatible plate reader using BioTek Synergy H1 (Agilent). Concentrations were calculated from the standard curve using a 5-parameter logistic fit, applying dilution factors to derive final values. Assays were accepted when the standard curve achieved R² ≥ 0.99 and QC recovery was within ±15% of the nominal value; samples outside the range were re-assayed following appropriate dilution and QC recovery was as expected. Qp was calculated using the formula: Qp (pg/cell/day) = Antibody titer (pg/mL) / (IVCD × Culture duration), where IVCD is the integral viable cell density. IVCD = ½ * (VCD at t₁ + VCD at t₂) * (t₂ - t₁) + IVCD at t₁.

### 2.11. Stability antibody production assay

Both pool and clones were expanded and serially passaged under routine cell maintenance until the passage P20, P40, or P60. Cells were maintained in CD FortiCHO™ medium under suspension conditions in unbaffled 125 mL Erlenmeyer shake flasks (30 mL working volume) at 37°C, 8% CO₂, and 125 rpm. Cells were subculture every 3–4 days to 0.3 × 10⁶ cells/mL, ensuring viability ≥ 90% at each split. The titer antibody assay was performed as mentioned previously in section 2.11. Supernatants were collected at day 7, 9, 12 and 14, at 125 rpm for 7 min, then transferred to 1.5 mL tubes (Millipore-Sigma, Cat#MTCC2000-500EA), and stored at −80°C.

### 2.12. Digital PCR (dPCR)

The insertion number was evaluated by transgene copy number per diploid genome and quantified by dPCR (QIAcuiy, Qiagen). gDNA was extracted from either pooled cells or isolated clones using the DNeasy Blood & Tissue Kit (Qiagen, Cat#69504). Samples were prepared from frozen pellets collected either during exponential growth for early passages between G0–G3 or during antibody production on days 7 and 14 for later G20, G40, G60.

Primers and probes were designed to specifically amplify the trastuzumab light-chain coding sequence of the transgene, with a probe FAM-labeled targeting the amplicon. For normalization, the Glucagon gene was used as the housekeeping reference with a HEX-labeled probe. Controls included gDNA from CHO-K1 wild type, used as negative control to verify transgene assay specificity and confirm the housekeeping copy number, a no-template using water was the negative control of the reaction, and a transgene expression plasmid was used as positive control. Copy number was reported as copies per diploid genome. The primers and probes sequences are listed in table 2.

### 2.13. Off-targets and on-target analysis after Cas-CLOVER editing

Genomic DNA from suspension CHO-K1 wild type (WT), 7G2 (GS5-SKO) clone, and CleanCut GS clone (GS-DKO) was sequenced by whole genome sequencing (WGS) as 151 bp paired-end reads using Illumina sequencing with a target of 60x coverage by SeqCenter, LLC (Pittsburgh, PA). The reads were aligned to the reference genome assembly CriGri_1.0 using Minimap2; the genome assembly was used for the other downstream analyses. For each gRNA sequence targeting the GS5 or GS1 loci respectively (Table 1), 23 bp off-target candidate sequences were identified using the CRISPOR web tool. The top 10 off-target candidate sites were selected for each gRNA sequence, giving a total of 40 sites based on the highest cutting frequency determination (CFD) off-target score. For off-target analysis, inspection intervals were constructed by padding each 23 bp off-target coordinate by ±500 bp to yield 40 off-target sites, each 1,023 bp in length. These intervals were used to extract locus-specific read subsets from whole-genome BAMs for each cell line.

The on-target analysis was performed to validate our previous genotype results and as internal control for our analysis by confirming the expected deletions from the edited GS KO clone cell lines. Read subsets were also extracted corresponding with the entire gene spans for Glul and LOC100689337.

Variant calling was done on these read subsets using FreeBayes v1.3.10. When variants (indels, SNP, or MPN) were detected within an on- or off-target interval, their displacement from the nominal target site was quantified as Distance = position target − position alternative, where the on-target position was defined as the mean of the outermost coordinates set by the paired left/right guides at the locus (rounded up to an integer) and the off-target position was defined as the mean of the two CRISPOR off-target coordinate endpoints (rounded to an integer). For on-target sites, only variants within 1000 bp were considered relevant. Variant origin was determined by sorting the variant calling outputs for all three cell line genomes (CHO-K1WT cells line, 7G2 and CleanCut GS clones) and identifying variant overlap. If a variant was present in all three genomes, the origin was considered Wildtype; if present only in 7G2 and CleanCut GS, origin was considered 7G2; if present only in CleanCut GS, origin was considered CleanCut GS. The interactive genome viewer (IGV) desktop application was used to visually validate variant origins and on-target deletions.

### 2.14. Data visualization and software

Data analysis was performed in GraphPad Prism v.10.0 and Microsoft Excel. Indel visualization and fragment analysis use PeakScanner v1.0. SnapGene was used for primer and construct design and Indel analysis, while ICE (Synthego) and Geneious Prime were used for sequence alignment and mutation calling. The software used for off-target and on-target analysis were as follows: Minimap2 v2.30-r1287, samtools v1.22.1, bedtools2 v2.31.1, gatk v4.6.2.0, FreeBayes v1.3.10, vcflib v1.0.3, and IGV v2.19.6.

## 3. Results

### 3.1. Cas-CLOVER achieves higher GS editing efficiency than Cas9 using Neon transfection system

Cas-CLOVER is a next-generation genome editing system that achieves high-efficiency cutting and indel formation. It functions as a dimeric nuclease, guided by paired left and right gRNAs that bring the two Clo051 nuclease domains into proximity, enabling precise cleavage at the intended target site. This design confers greater specificity than conventional Cas9 and forms the basis for our comparison of editing efficiencies in CHO-K1 cells (Figure 1A). Neon transfection efficiency was first assessed using a GFP-expressing mRNA as control, achieving 98% GFP-positive cells and 89% viability post-transfection (Figure S2A, B). We then optimized the gRNA to Cas-CLOVER mRNA ratio by testing one gRNA pair targeting GS5 and another targeting GS1. At an optimal 2:1 gRNA:mRNA ratio, editing efficiencies measured by T7 endonuclease assay at 72 h post-transfection reached 92% for GS5 and 98% for GS1 (Figure S2 C-F). To compare editing efficacy using Neon transfection, Cas-CLOVER and Cas9 mRNA were transfected with either paired gRNAs (for Cas-CLOVER) or with individual L- or R-gRNAs (for Cas9), maintaining the 2:1 ratio (gRNAs:mRNA). For GS5, Cas-CLOVER achieved 84% editing efficiency, while Cas9 using L- or R-gRNAs reached 57.4% and 30.4%, respectively (Figure 1B, D). For GS1, Cas-CLOVER showed a significant difference of 74% editing efficiency compared to 62% and 54.4% for Cas9 with L- and R-gRNAs, respectively (Figure 1C, E).

**Figure. 1.**
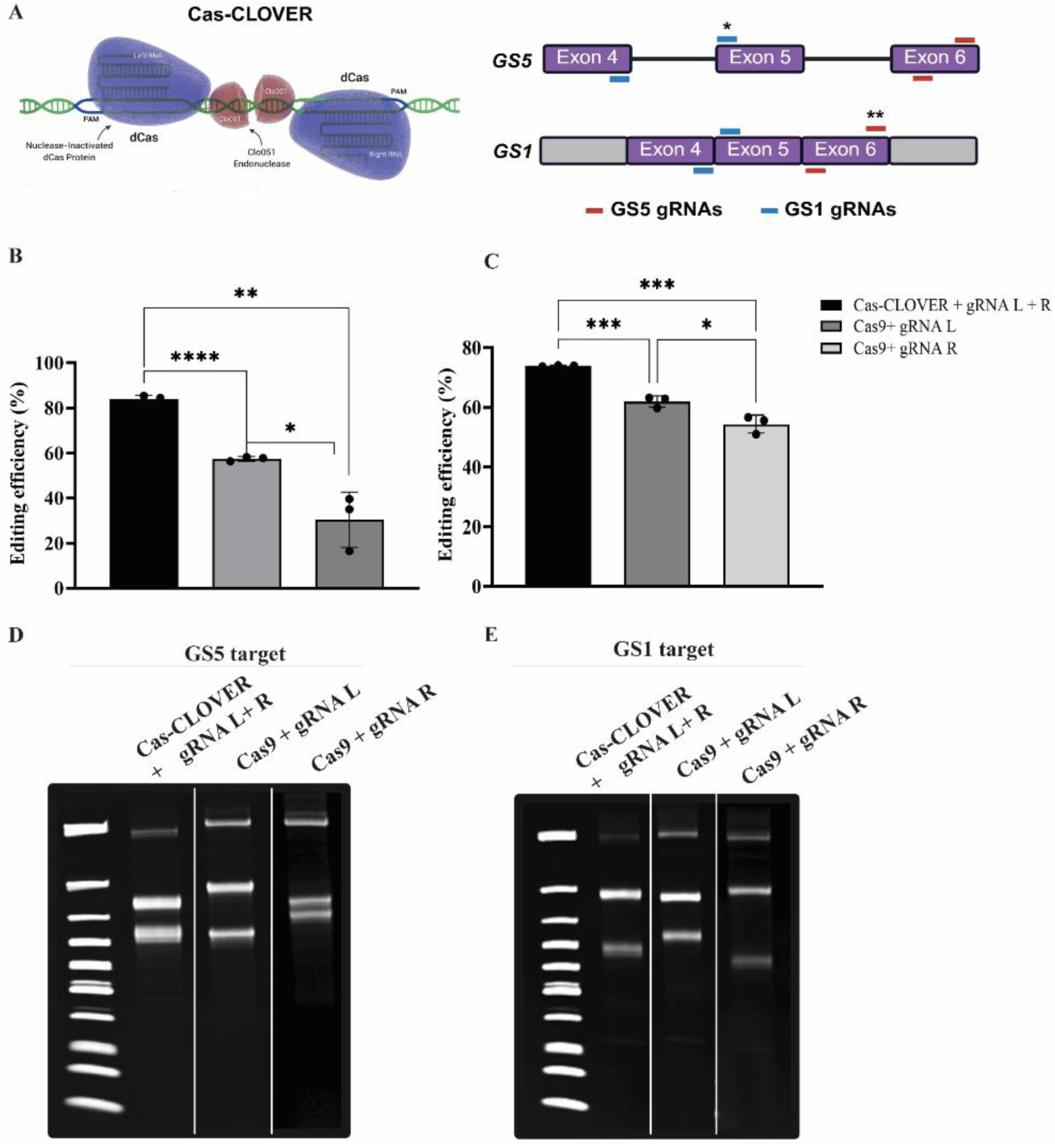
Efficient disruption of GS in suspension CHO-K1 cells using Cas-CLOVER. **(A)** Schematic representation of the Cas-CLOVER complex with gRNAs targeting the GS5 and GS1 genomic regions. One gRNA pair was designed to target exon 6 of the GS5 gene, and another gRNA pair was designed for exon 1 of the GS1 gene at the junction homologous to chromosome 5 (exons 4–5). Single (*) and double (**) mismatches indicate gRNA binding variations at the target site. **(B, C)** Editing efficiency quantification of Cas-CLOVER targeting GS5 and GS1 compared with Cas9, respectively. **(D, E)** Editing efficiency was assessed using the T7EI mismatch assay with CHO-K1 cells that harvested 72 h post-transfection. Representative gels show the expected band patterns for each condition. For GS5 editing, the unedited band was 752 bp. Cas-CLOVER generated 413 bp and 339 bp fragments, while Cas9 produced 752 bp (unedited), 438 bp and 314 bp (gRNA-L), and 388 bp and 364 bp (gRNA-R). For GS1 editing, the unedited band was 796 bp. Cas-CLOVER generated 487 bp and 309 bp fragments, while Cas9 produced 796 bp (unedited), 462 bp and 334 bp (gRNA-L), and 512 bp and 284 bp (gRNA-R). Experiments were performed in biological triplicates, and error bars represent mean ± SD. Statistical significance was determined using Student’s *t*-test, with *P* < 0.05 considered significant.

### 3.2. Cas-CLOVER Mediated GS5-SKO and GS-DKO Knockouts

To establish an edited CHO platform suitable for commercial CLD and GMP-scale manufacturing, adherent CHO-K1 cells were first adapted to serum-free suspension culture in CD FortiCHO medium, achieving >90% viability and complete adaptation after several passages. Following this, cells were transitioned to glutamine-free conditions over 7 days. Using the adapted platform, we generated GS5-SKO cells and GS-DKO cells at both GS5 and GS1 loci through a sequential transfection and selection strategy (Figure S3). To generate the GS5-SKO cell line, cells were transfected with Cas-CLOVER targeting exon 6 of the GS5 locus by nucleofector giving an editing efficiency of 47% (Figure S4).

SCC was carried out using multiple 96-well plates to ensure comprehensive recovery of edited clones. Putative clones were systematically expanded, screened by PCR for GS knockout edits, and further confirmed by Sanger sequencing to validate genotypes at high resolution. This approach identified multiple clones with biallelic, out-of-frame indels, most falling within the 20–30 bp deletion range, including one with a 248 bp deletion, 90.4% of GS5 editing was identified in a range of 0 to −45 bp Net deletion + insertion in GS5-SKO clones (Table 3, Figure 2A). After monoclonality confirmation (Table 3), several top clones (6E7, 9E11, 8H7, 7G2, 7C4, and 6C9) were selected for growth kinetic evaluation, with sequence alignment of their GS5 genotypes serving as a key criterion for choosing the final candidate clone (Figure 2B, Table 4).

**Figure 2.**
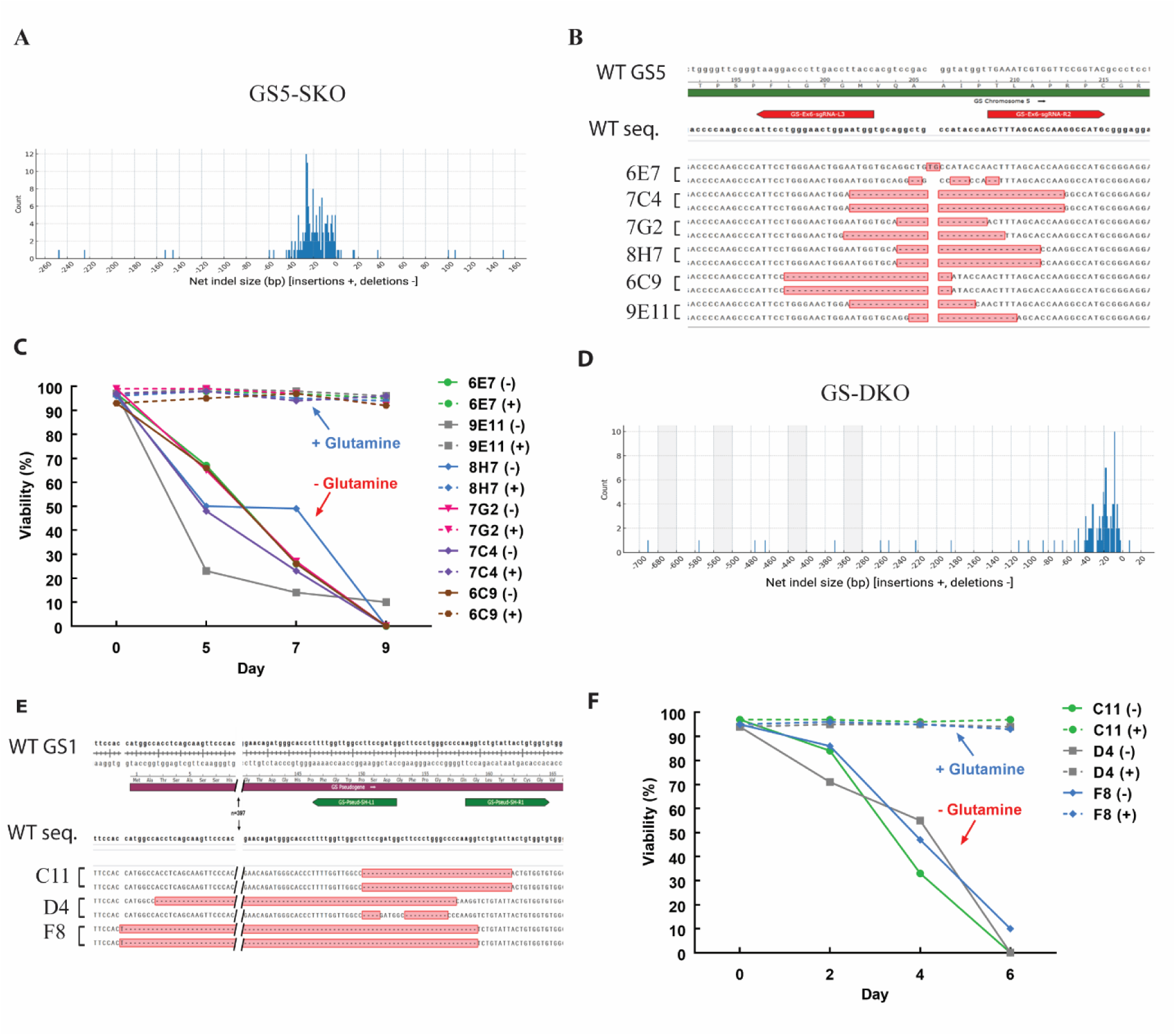
Cas-CLOVER -mediated knockout of GS5 and GS1 genes in CHO cell lines and their impact on cell viability. **(A)** Indels frequency after SCC genotyping in GS5-SKO CHO cell line. **(B)** Sequence alignment of the GS5 locus with reference WT CHO cells sequence and GS5-SKO clones (6E7, 7C4, 7G2, 8H7, 6C9, and 9E11) for genotype screening. (**C)** Cell viability of GS5-SKO clones cultured in CHO medium with (+) or without (–) L-glutamine over 9 days. **(D)** Indels frequency after SCC genotyping in GS-DKO CHO cell line. **(E)** Sequence alignment of the GS1 locus with reference WT CHO cells sequence and GS-DKO clones (C11, D4, and F8) for genotype screening. **(F)** Cell viability of GS-DKO clones cultured in medium with (+) or without (–) L-glutamine over 6 days.

**Table 3.** Total clones number summarization to select the top edited clone for GS5-SKO or GS-DKO CHO cell lines.

| Steps for clone screening | GS5-SKO | GS-DKO |
| --- | --- | --- |
| 96-well lates | 3 | 8 |
| Seeding Single Cells (VIPS) | 231 | 558 |
| Clones with Confluence $\geq 15\%$ | 129 | 282 |
| Consolidated Clones | 108 | 243 |
| Clones Screened by PCR | 108 | 83 |
| Sequence-Confirmed Clones | 64 | 78 |
| Clones with Out-of-Frame Indel | 20 | 39 |
| Clone after Monoclonality | 6 | 3 |
| Final top clone | 7G2 | F8 |

**Table 4.** Indel profiles of GS knockout CHO-K1 cell lines.

| Genotype | Clone | indels |
| --- | --- | --- |
| SKO | 6E7 | 2bp ins, 7bp del+ 6bp ins |
|  | 7C4 | 34bp, 34bp del |
|  | 7G2 | 25bp, 13bp del |
|  | 8H7 | 22bp, 22bp del |
|  | 6C9 | 26bp, 26bp del |
|  | 9E11 | 19bp, 16bp del |
| DKO | C11 | 35bp del |
|  | D4 | 465bp, 14bp del + 6 bp ins |
|  | F8 | 477bp del + 1 bp ins |

*Note:* Summary of insertion and deletion (indel) mutations identified GS5-SKO and GS-DKO CHO-K1 clones generated using Cas-CLOVER. Indel sizes are shown in base pairs (bp).

A glutamine sensitivity assay was established to evaluate clone dependence on endogenous GS activity by comparing growth in the presence or absence of L-glutamine. Viability and total cell number were measured on days 5, 7, and 9, confirming complete loss of proliferation under glutamine-free conditions. In contrast, robust growth was observed in the presence of glutamine, demonstrating that exogenous supplementation is required for proliferation (Figure 2C). Based on this sensitivity profile, clone 7G2 was selected for further characterization. The growth kinetics of GS5-SKO clone 7G2 were monitored over 14 days under standard culture conditions. VCD was measured every two days, and cultures were re-seeded at 0.3 × 10^6^ cells/mL after each count to maintain exponential growth. VCD measurements showed consistent expansion with average doubling times of 24 h, confirming stable viability and proliferation throughout the evaluation period (Figure S5A).

To generate GS-DKO cells, the 7G2 clone was nucleofected with Cas-CLOVER targeting exon 1 of the GS1 locus corresponding to the homologous region on chromosome 5 (exon 4 and 5). Giving an editing efficiency of 46.3% (Figure S4B). Despite a sequence similarity of 96.7% between GS5 and GS1, Cas-CLOVER’s requirement for nuclease dimerization enabled precise editing of GS1 while leaving GS5 unmodified (Figure 3), as the intervening intron at the GS5 locus prevented Clo051 dimerization.

**Figure 3.**
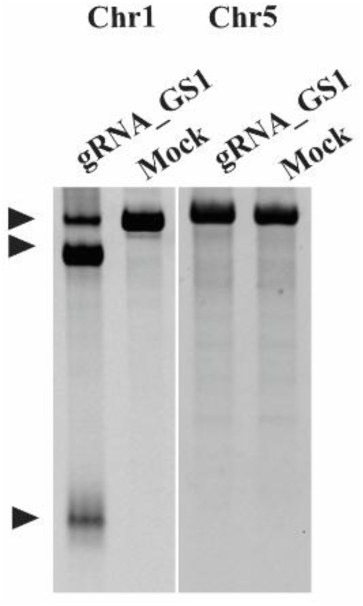
Representative gel analysis of RNA-guided editing strategies targeting GS1 and the GS5-SKO (7G2) clone. A distinct cleaved band pattern was observed when gRNAs recognized target sequences on GS1, yielding fragments of 618 bp and 152 bp from the unedited 770 bp band. No cleavage was detected when guide pairs bound without sufficient proximity (GS1 guide on GS5).

SCC across eight 96-well plates yielded 558 colonies, of which 282 reached confluence and 243 were expanded for further analysis. GS-DKO clones were identified by PCR and confirmed by Sanger sequencing (Table 2). Thirty-nine clones carried out-of-frame indels, with a broad distribution of deletion sizes similar to those observed at GS5, 85.8% of GS1 editing was identified in a range of 0 to −50 bp Net deletion + insertion in GS-DKO clones (Figure 2D). After monoclonality confirmation, six top clones with indels ranging from 1 bp to 477 bp were selected for further characterization (Table 4, Figure 2E).

Clones C11, D4, and F8 were selected for growth kinetic analysis to identify the top candidate. A glutamine sensitivity assay was performed by measuring viability and total cell number; however, due to the rapid loss of viability in GS-DKO clones, measurements were taken on days 2, 4, and 6. This confirmed an enhanced loss of proliferation caused by the absence of endogenous GS activity (Figure 2F). Based on this sensitivity profile, clone F8 was selected for further characterization. VCD measurements showed steady expansion with an average doubling time of 21.3 h, confirming that clone F8 maintained stable viability and proliferation throughout the evaluation period. This clone was designated as the CleanCut GS platform (Figure S5B). In addition, GS gene expression was absent in CleanCut GS clone quantified by qPCR as shown in figure S5C.

### 3.3. CleanCut GS Cells pool and clone Enable High-Titer Trastuzumab Production

Figure S6 provides an overview of the comparison study evaluating the performance of GS5-SKO (clone 7G2) versus CleanCut GS cells. Both cell lines were transfected with a transposon vector encoding trastuzumab with heavy and light chains, along with glutamine synthetase for glutamine selection and Harbor-IN transposase mRNA. Stable pools were analyzed in 24-deep well plates for antibody titer and Qp, with top clones isolated from the CleanCut GS pool expressing trastuzumab (CleanCut GS-TZ) subjected to further analysis. The GS5-SKO pool achieved an antibody titer of 1.65 g/L and a Qp of 55.2 pg/cell/day, whereas the CleanCut GS-TZ pool reached a peak titer of 4.21 g/L and a Qp of 83.9 pg/cell/day (Figure 4A, B, respectively). Figure 4C outlines the cell-specific production of trastuzumab over the course of 14 days in the CleanCut GS-TZ pool. The CleanCut GS-TZ pool also demonstrated robust proliferation, reaching a peak VCD of 12.6 × 10⁶ cells/mL on day 7 while maintaining >96% viability, highlighting its adaptability to cyclic feeding conditions even at late culture stages (Figure 4D).

**Figure 4.**
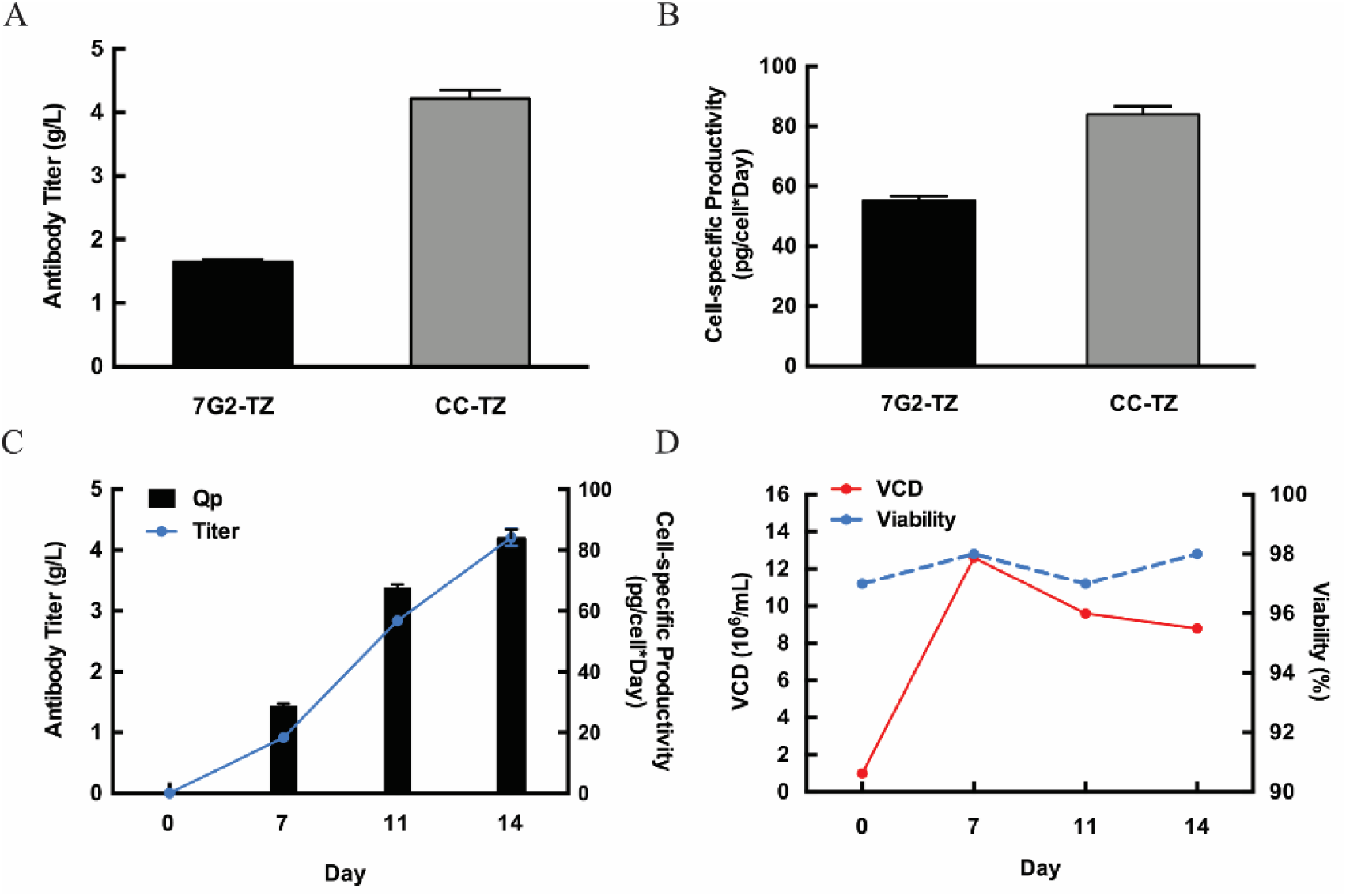
GS5-SKO and GS-DKO Trastuzumab productivity comparison. **(A)** Titer comparison for GS5-SKO-TZ and GS-DKO-TZ on day 14. **(B)** Specific productivity (Qp) comparison for GS5-SKO-TZ and DKO-TZ on day 14. **(C)** Titer and specific productivity (Qp) of the CleanCut GS-TZ (CC-TZ) pool at different culture days. **(D)** Viable cell density (VCD) and viability of the CC-TZ pool at different culture days. Experiments were performed in triplicates, and error bars represent mean ± SD.

Given this improvement, single-cell cloning was performed on the CleanCut GS-TZ, yielding two top clones, 5G2-TZ and 31H3-TZ. Clone 5G2 reached a peak VCD of 9 × 10⁶ cells/mL on day 14 while maintaining >96% viability through day 13 and 94% at harvest (Figure 5A). Antibody titers increased from 3.9 g/L on day 12 to 5.6 g/L on day 14, with Qp rising from 80.2 to 109.4 pg/cell/day, surpassing pool productivity (Figure 5B). To assess earlier production kinetics, clone 31H3-TZ was monitored at mid-culture timepoints. This clone achieved a VCD of 9.4 × 10⁶ cells/mL and 8.6 × 10⁶ cells/mL on days 7 and 14 respectively, with >96% viability throughout, while titers increased from 1.5 g/L on day 7 to 5.3 g/L on day 14. Correspondingly, Qp rose from 55.3 pg/cell/day to 108.4 pg/cell/day by harvest (Figure S7A, B). These findings position CleanCut GS as a ready-to-use, high-productivity CHO platform, well-suited for GMP-ready biologics manufacturing.

**Figure 5.**
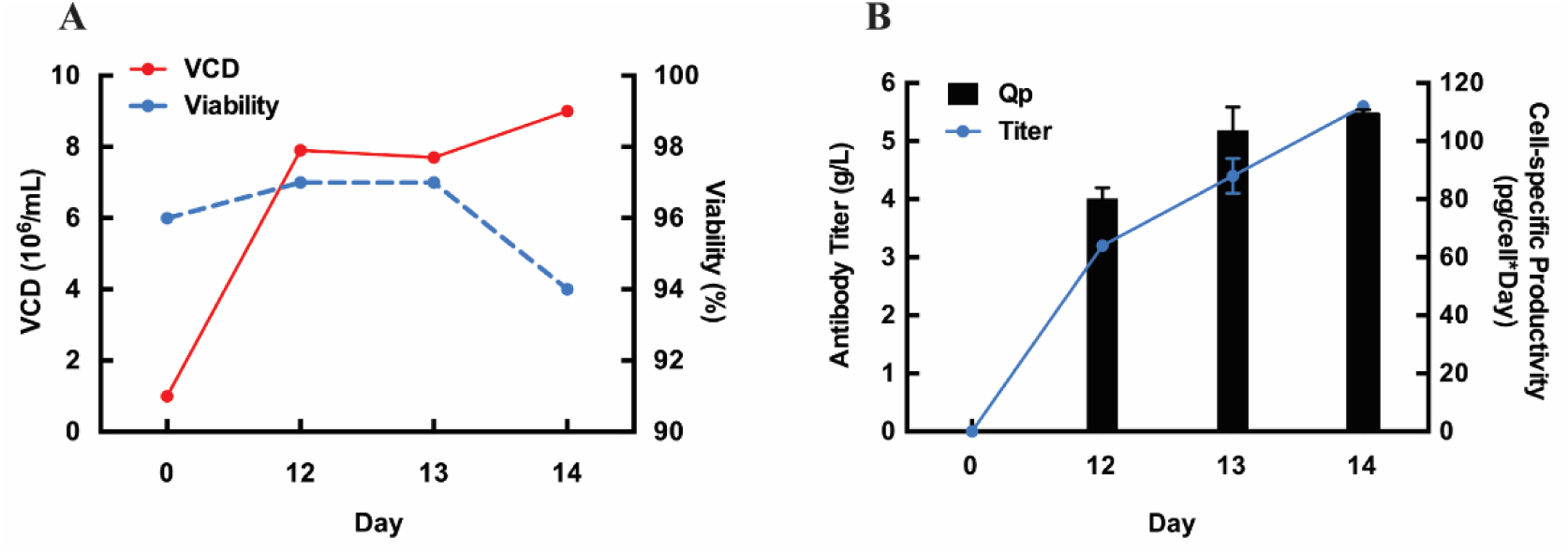
Characterization of GS-DKO CHO cell expressing Trastuzumab. **(A)** Viable cell density (VCD) and viability of the 5G2-TZ clone for a 14-day culture. **(B)** Titer and specific productivity (Qp) of 5G2-TZ for 14 days culture. Experiments were performed in triplicates, and error bars represent mean ± SD.

### 3.4. A stability analysis shows titer production consistency in CleanCut GS-TZ pool and clone over 60 generations

In order to evaluate the long-term stability of insertions by Harbor-IN transposase, we quantified titer, Qp, and VCD, in the CleanCut GS-TZ pool and in the isolated 5G2-TZ clone across 60 generations. Within each fed-batch, both pool and clone showed an expected increase in titer and Qp overtime on days 7, 9, 12, and 14. For each cell line, we observed similar results across passages 20, 40 and 60 (Figure 6). VCD remained stable for both CleanCut-TZ pool and 5G2-TZ clone (Figure S8). On day 14, the CleanCut GS-TZ pool achieved a titer of 4.3 g/L and a Qp of 93.39 at generation 20, 4.87 g/L and 70.49 at generation 40, and 4.59 g/L and 76.72 pg/cell/day at generation 60 (Figure 6A, C), with corresponding VCD values of 7.5 × 10^6^ cells/mL, 13.6 × 10^6^ cells/mL, and 10.8 × 10^6^ cells/mL (Figure S8A). These data are consistent with the initial characterization that we consider generation 0 of the pool for both titer and Qp, supporting its phenotypic stability over extended passaging.

**Figure 6.**
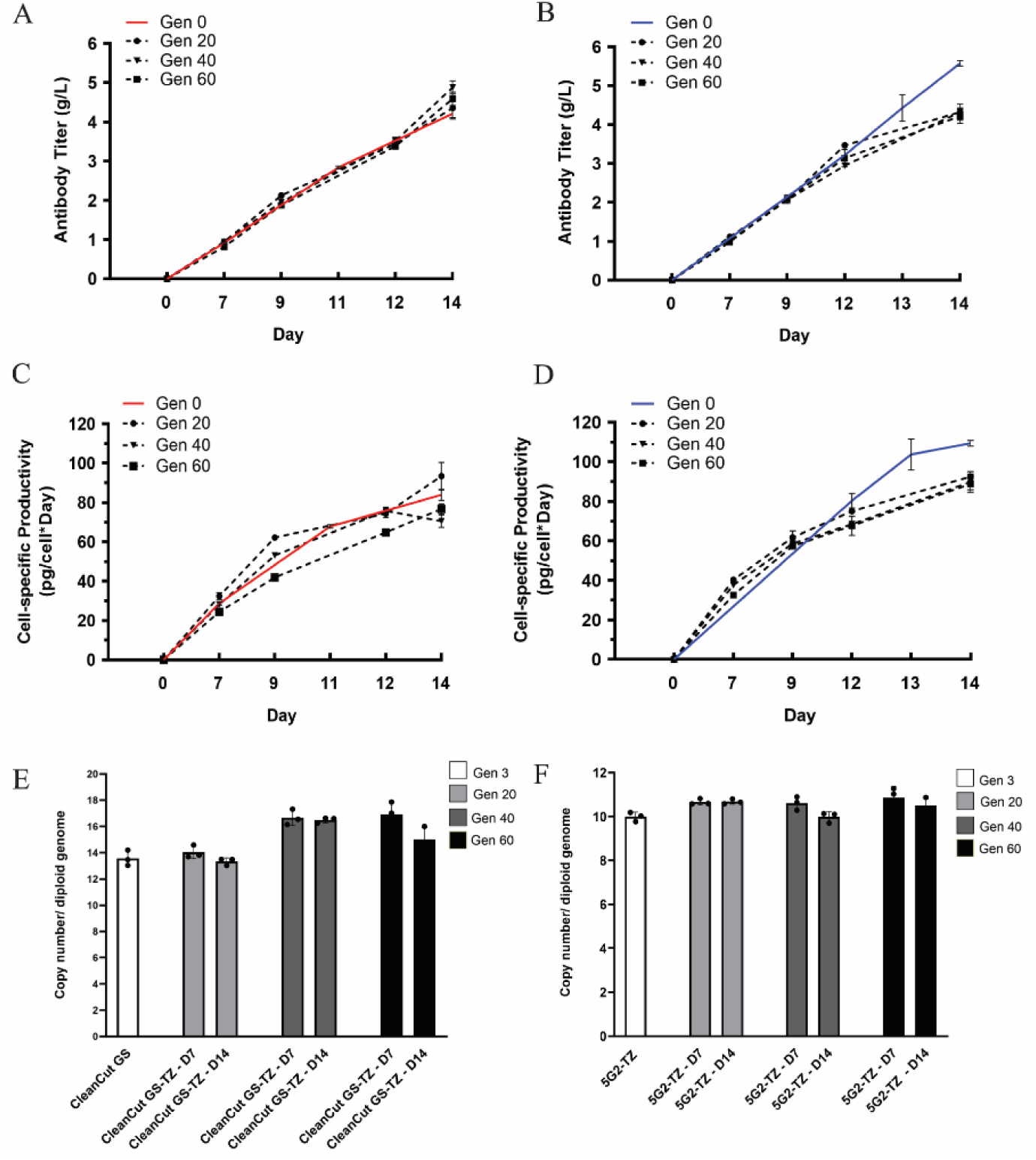
Stability analysis of CleanCut GS-TZ pool and 5G2-TZ clone over 60 generations. **(A, B)** Titer and **(C, D)** Specific productivity (Qp) quantification was evaluated in CleanCut GS-TZ pool and 5G2-TZ clone over 20, 40 and 60 generations and compared with Generation 0 for 14 days. **(E, F)** Genomic copy number per diploid cell was quantified using the trastuzumab light-chain sequence previously integrated via the Harbor-IN transposase system in CleanCut-TZ and 5G2-TZ. Data is presented as mean ± SD from three independent replicates.

We also evaluated the isolated clone, which showed consistency in the titer and Qp rising from day 7 to day 14 over the generations. Specifically, on day 14 the clone reached a titer of 4.31 g/L and a Qp of 92.4 at generation 20, 4.3 g/L and 89.8 at generation 40, and 4.21 g/L and 88.77 at generation 60 (Figure 6B, D), with VCD values of 6.9 × 10^6^ cells/mL, 8.22 × 10^6^ cells/mL, and 7.07 × 10^6^ cells/mL, respectively (Figure S8B). The clone exhibited consistent expressions across all time points through day 12, with only a modest reduction of approximately 20% observed on day 14. This decrease remained within the predefined acceptance threshold of ±30% after 60 generations and was therefore considered acceptable.

In order to corroborate stability across generations, transgene insertion numbers were measured in the CleanCut GS-TZ pool and the 5G2-TZ clone at G3, G20, G40, and G60 during titer production on days 7 and 14. Trastuzumab was used as the target for genomic copy-number quantification, normalized to the glucagon housekeeping gene. Insertion number per cell was correlated with the copy number per diploid genome. In the CleanCut GS-TZ pool, 14 insertions per cell were observed at G3 and G20 on both day 7 and day 14, increasing to 16 insertions at G40 and G60. At G60, the value diminished on day 14 (Figure 6E). Under the same conditions, the 5G2-TZ clone maintained 10 insertions per cell across all generations and time points (Figure 6F), indicating greater clonal consistency than the pooled population measured via dPCR.

### 3.5. Protein-level characterization and glycan profile of trastuzumab secreted by GS-DKO-TZ pool

The trastuzumab antibody produced by the GS-DKO-TZ pool was purified and confirmed the presence of both light and heavy chains by SDS-PAGE (Figure S9A). In figure S9B, we further analyzed the stochiometric of the light chain and heavy chain ratio using a reduced-protein liquid chromatographic profile. The light chain appears at 4.66 min, while the heavy chain elutes at 5.14 min. Because the heavy chain is approximately twice the molecular weight of the light chain, we expected a 1:2 light chain: heavy chain ratio in integrated peak areas, the heavy chain peak displays roughly double the signal of the light chain peak. To further strengthen the product characterization, we evaluated the glycan profiles of CleanCut GS-TZ pool. The analysis revealed a predominant abundance of G0F and G1F glycans on the heavy chain, representing 61.25% and 7.7%, respectively, whereas no detectable glycan residues were observed on the light chain by reduced LC-MS (Figure S9C, D).

### 3.6. Cas-CLOVER edits GS1 and GS5 with none detected off-targets

To evaluate the precision of Cas-CLOVER editing in the GS5-SKO and GS-DKO clones, off-target and on-target events were analyzed using Illumina whole genome sequencing reads. A CHO-K1 WT was included in the analysis to differentiate Cas-CLOVER off-target events from pre-existing variation between the parental WT genome and the reference genome. Table S1 contains the 40 genomic off-target sites investigated in GS5 and GS1 loci. Variants calling in 40 off-target sites, representing 10 off-target sites per gRNA used for Cas-CLOVER editing. Aligning the unmerged paired-end reads from CHO-K1 WT, 7G2, and CleanCut GS clones to the CriCri_1.0 genome produced mean genome-wide read depths of 155.307x, 125.757x, and 140.514x, respectively. The analysis revealed a total of 31 variants (classified as SNPs, MNPs and indels) presented in the three samples. We identified variants in the off-target sites in an allelic frequency between 0.5-1, these variants were found in the CHO-K1 WT, indicating that they are originally from the parental cells (Table S2) and not from undesired cleavages by Cas-CLOVER.

We evaluated the on-target site by aligning the corresponding GS5 or GS1 loci with CHO-K1 WT. On-target analysis revealed 13bp and 25bp deletions in the GS5 site originating in 7G2 (Figure 7A, Table S3). The finding indicated one SNP 3bp from the on-target site in GS1 (C to G) originating in CleanCut GS. Three additional variants were identified as originating in CHO-K1 WT (Figure 7B, Table S3). The expected large deletion in GS1 was verified by IGV as a large span of reads missing from the on-target site in CleanCut GS (Figure 7B). All on- and off-target sites were also visualized with IGV, confirming that variant calling was accurate and no prominent indels were missed by variant calling.

**Figure 7.**
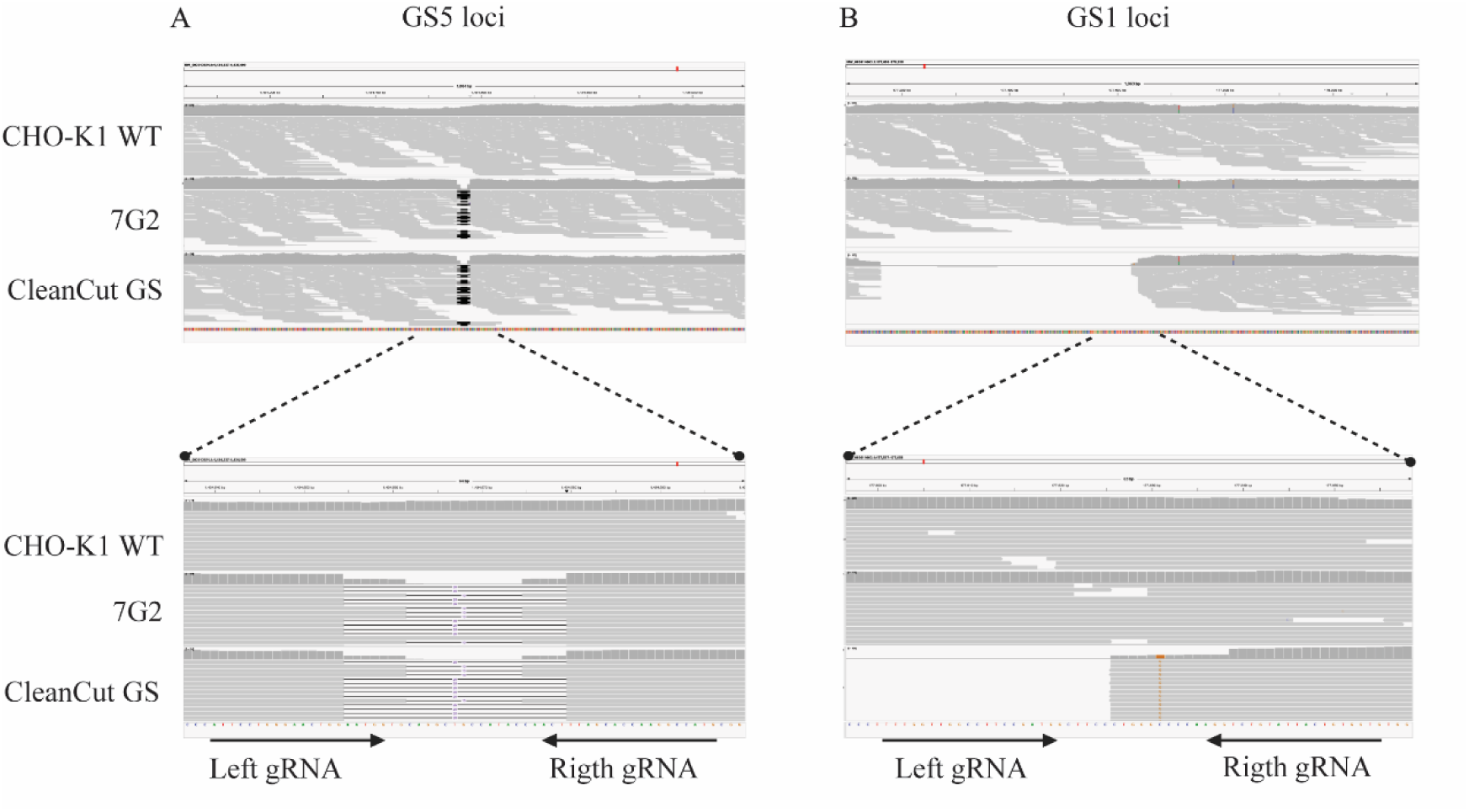
Representative images of GS KO CHO cells lines genotype verified by WGS and compared with the CHO-K1 WT. **A.** Two deletions were identified in the 7G2 and CleanCut GS in the GS5 loci (top). The regions show two deletions of 13bp and 25bp (bottom). **B.** A large deletion was identified only in CleanCut GS clone in the GS1 loci. The regions show a fragment of the reads corresponding with the large deletion (bottom). The left and right borders show the PAM sites for the on-target, arrows represent the left and right gRNAs.

In addition, the on-target analysis in GS1 loci revealed a SNP only in one site in CleanCut which was a SNP (C to G) of 3 bp from the GS1 on-target site, however, it was not identified in the CHO-K1 WT and 7G2 indicating this genotype may associated with Cas-CLOVER editing events. Finally, two SNPs in different locations were identified in the three samples which suggest this alteration is originally from the CHO-K1 WT parental cell line.

## 4. Discussion

CHO cells remain the workhorse for biopharmaceutical production, but efficient and precise genome editing tools can accelerate CLD and improve the manufacturing of complex new modalities. While CRISPR/Cas9 is widely adopted in research, its application in commercial CLD and GMP settings is limited by elevated off-target activity, leading to chromosomal instability, and a complex intellectual property landscape that increases costs and slows adoption ^22^. In this study, we evaluated Cas-CLOVER as a genome editing solution uniquely suited for industrial CLD applications to overcome these barriers. Our goals were to test Cas-CLOVER efficiency in suspension-adapted CHO-K1 cells compared to the conventional CRISPR/Cas9, assess its specificity by sequentially targeting two highly homologous GS loci, and establish a gene-edited production platform, the CleanCut GS, that could be validated with the Harbor-IN transposase system for stable antibody expression. By using an optimized protocol together with the GS-DKO cell line for stable antibody production, we achieved higher titer yields in pools than in isolated clones.

Cas-CLOVER demonstrated consistently higher editing efficiency than Cas9 with Neon transfection showing greater than 74% editing observed across GS5 and GS1 loci (Figure 1). Importantly, the Clo51 endonuclease dimerization requirement of Cas-CLOVER not only reduces off-target activity but also confers unique specificity and target design benefits. In this study, targeting the exon junction of GS1 provided precise editing while leaving the homologous GS5 locus unmodified, highlighting the system’s ability to discriminate between highly similar sequences (Figure 3). Finally, our findings expand upon the cell types previously demonstrating Cas-CLOVER’s high editing efficiencies in HEK cells and T cells ^23, 18^.

Beyond this case, the dimeric design of Cas-CLOVER also facilitates homologous recombination donor vector strategies: plasmids are not cleaved by Cas-CLOVER due to the intervening donor insert and lack of dimerization, eliminating the need for engineering silent PAM mutations that are often required with Cas9. Cas-CLOVER also generates larger deletions in CHO cells, as observed after genotyping confirmation (Figure 2), confirming an earlier finding in HEK293 cells showing also larger deletions ^23^. One of our previous studies described the use of Cas-CLOVER to KO the *STAT1* gene in HEK293 cells. They measured the expression of the viral proteins VP1, VP2, and VP3, which constitute the AAV capsid, in isolated STAT1 KO clones and quantified vector genome titers in comparison with wild-type HEK293 cells ^23^. Additionally, another study examined the editing efficiency of Cas-CLOVER in T cells and iPSC when combined with a transposon system for CAR-T cell therapy ^18^. Together, these studies demonstrate the potential of Cas-CLOVER for different cell editing and highlight its applicability to biopharmaceutical and cell therapy applications.

To evaluate the high precision of Cas-CLOVER reported by Madison et al. (2022) ¹⁸, we performed WGS analysis of predicted off-target sites based on the Left and Right gRNAs in GS5 and GS1 loci, detecting off-target events by Bayesian variant detection. Indeed, the analysis of the 40 most likely off-target guide binding sites, including one site with 100% sequence identity to the gRNAs, revealed none detected off-target effects. We confirmed that there was no major difference between the three samples by IGV, with an average of reads of 150 bp per sequence, and none detected off-target events were found within 500 bases of the off-target sites on the left or right side. Therefore, while this analysis was restricted to candidate sites and did not include unbiased detection or structural variant and translocation screening, these results provide evidence that no off-target events were detected in the 40 sequences predicted by the CRISPOR tool. Additional genome-wide analyses would be required to confirm the off-target event in the CHO genome.

Recent studies have shown several advantages of Cas-CLOVER over Cas9 and earlier gene-editing tools such as the TALEN and ZFN system including editing efficiency, cost-effectiveness, time-consuming protocols and intellectual property considerations and off-target frequency ^24,25^. Among the advantages Cas-CLOVER produces between 3 to 5 bp 5′ overhangs, which are more conducive to precise ligation or targeted knock-ins and often reduce undesired indels, whereas Cas9 generates blunt ends that are more prone to error-prone repair ^26^. In addition, Cas-CLOVER pairs efficiently with PiggyBac transposons to insert large DNA cargos. Cas-CLOVER is also associated with a lower risk of chromosomal rearrangements ^18^. Because double-strand breaks occur only when both halves of the dimer are correctly positioned, the system results in a lower chromosomal translocation, lower genotoxicity, and fewer unintended breaks. In addition, Cas-CLOVER requires two guide RNAs, which provides an additional layer of specificity by avoiding sites near off-target homologous sequences ^18^. As Cas-CLOVER is a relatively new gene-editing tool, it is important to evaluate its performance across additional cell types to confirm its efficiency more broadly. Since cell therapy includes a variety of primary cells such as hematopoietic stem cells, mesenchymal stem cells, and induced pluripotent stem cells it would be valuable to assess the editing efficiency of Cas-CLOVER in these cell populations as well.

In this study, we evaluated for the first time the editing efficiency and CHO GS DKO cells characterization using Cas-CLOVER for biopharmaceutical applications. The GS selection system enables rapid generation of antibody producing CHO pools and clones in about two weeks, with or without MSX. A study showed that monoclonal antibody production increased in two top GS KO clones in absence of MSX compared with the conditions incubated with MSX. The authors found that compared with CHO-K1 cell pools cultured with 25 µM MSX, GS knockout pools grown without MSX did not show higher monoclonal antibody production, supporting that the GS selection system offers a shorter timeline for establishing the cell line and the ability to avoid MSX during both cell line generation and routine culture ^27^. The GS gene at the GS5 locus is among the most frequently targeted sites in CHO cells, as reported using different gene editing tools such as ZFN and Cas9 technology ^8,28,29^ and a pseudo GS gene was recently disrupted to generate a GS-DKO CHO cell using CRISPR/Cpf1 ^11^.

Based on the GS-DKO reported on the literature, in this study, we edited the GS5 and GS1 loci to generate the GS5-SKO and GS-DKO pools and screened 110 and 83 clones for GS5 and 1 gene disruption, respectively (Table 3). In order to increase the probability of obtaining SCCs, we used VIPS as shown in other studies ^30^. After confirming monoclonality, biallelic GS5 and 1 KO clones were expanded to evaluate glutamine dependence. In the absence of glutamine, GS5-SKO clones were eliminated by day 9 and GS-DKO clones by day 6, supporting GS-DKO as a more effective selection approach, as shown in Figure 2. The genotyping revealed that CleanCut GS has a deletion of 25 bp and 13 bp in GS5 and a 477 bp deletion + 1bp insertion in GS1 confirming the disruption of both GS genes (Figure 2, Figure 7).

In this study, Harbor-IN transposase enabled the creation of stable trastuzumab-producing pools derived in both GS-SKO and CleanCut GS backgrounds. The optimized protocol used in this study fed daily the cells with a 2% (v/v) HyClone CellBoost™ supplements. On day 4, the temperature was downshift from 37 ° to 32 °C. Zhu et al, demonstrated that the temperature downshift from 37 to 33 °C on day 5, led to a higher cellular viability and increased antibody titer by 48% and 28% in high-producing and how-producing CHO cell lines, and reduced charge heterogeneity and molecular size heterogeneity ^31^. Feeding daily the cells prevent nutrients depletion such as glucose and other necessary amino acids ^31^. The GS-SKO pool achieved a titer of 1.65 g/L with a Qₚ of 55.2, whereas the CleanCut GS pool reached 4.21 g/L with a Qₚ of 83.9, representing a substantial improvement in productivity (Figure 4). The significant difference between both cell lines showed that the GS-DKO expressing TZ pool improved the titer and Qp on day 14. Based on these results, we decided to evaluate the antibody production in two isolated clones derived from CleanCut GS-TZ pool. The SCCs from this pool yielded top clones such as 5G2 and 31H3, which surpassed pool-level productivity with a titer of 5.6 and 5.3 g/L and a Qp measurements of 109.4 and 108.4 pg/cell/day, respectively (Figure 5, Figure S5), demonstrating the robustness of the CleanCut GS platform for biomanufacturing applications.

Over the years, several studies have evaluated stable transfection strategies for antibody production using cell pools rather than isolated clones, demonstrating that pool-based approaches can substantially shorten development timelines compared to traditional clone screening ^21, 32, 33^. New strategies for stable expression in CHO cells using a combination of transposases were recently evaluated in terms of copy number integration and their potential impact on antibody production ^34^. Rajendra et al. (2016) demonstrated that the Super piggyBac transposase system enabled CHO pools to produce antibodies after rapid selection (7–10 days), reaching titers of 2.8– 7.6 g/L and a Qp of 25–70 by day 14 in a 24-deep-well plate format. These results show that cell pools can achieve titers comparable to clonal lines and are readily scalable for production ^21^.

In this current study, the stability analysis for antibody production showed that the CleanCut GS-TZ pool-maintained titers and Qp in line with baseline measurements across 60 generations (Figure 6). Although the 5G2-TZ clone showed a slight decrease in titer and Qp over successive generations compared with the initial passages, this reduction remained within the industry-accepted stability threshold, typically defined as maintaining ≥70% of the original titer (i.e., ≤30% decrease), and was therefore considered stable ^35^.

The long-term stability of antibody production observed in the CleanCut GS-TZ pool supports the potential of both the CleanCut GS cell line and the Harbor-IN transposase for large-scale biopharmaceutical manufacturing, including production of IND-enabling toxicology and Phase I clinical trial material. In parallel, single-cell cloning can be performed to isolate and characterize top-producing clones for use in later-phase clinical material and commercialization. The insertions number per cell showed consistency in G20, 40 and 60 compared with G3 in both CleanCut GS-TZ and G2-TZ, confirming the transgene integrity across the generations of the characterized cell lines in this study. Recent studies evaluated the correlation between protein production and gene copy numbers, focusing on genetic instability due to the loss of transgenes across time. Qian et al., evaluated the clone-specific genetic instability that can result in an unstable antibody production, in which copy number was detected by qPCR on Day 8. The authors showed that light- and heavy-chain transgene copy numbers were stable from passages P12 to P21 and start to decline at P25 and dropped markedly by P37 together with the protein productivity ^36^. Dhiman et al., used 13 Horizon CHO cell lines classified in low (<3), medium (4-15) and high (>15) copies number to evaluate how the titer can diminish from passage 1 to 10 in terms of titer production based on the genomic, transcriptomic and epigenetic characteristics. They observed that < 25% of the titer was reduced in the group of stable expression with low and medium transgene copy number between P1 and P10 ^37^.

The use of stable pools offers a major advantage over the clones by significantly reducing development time and cost, enabling scale-up and translational manufacturing within weeks rather than the typical 9–12 months required for clonal selection and process development.

Our results can be compared with the recent report by Srila et al. (2023), which also investigated GS5 and GS1 double knockouts in CHO cells. Both studies demonstrated that targeting both GS loci improves selection stringency and enhances productivity ^11^. However, the CleanCut GS platform achieved substantially greater titers and cell-specific productivity. Several factors contributed to this difference. First, the use of Cas-CLOVER minimized off-target activity, likely enabling cleaner editing than Cas9/Cpf1-based approaches, and improving our cell platform development. Second, our workflow incorporated more rigorous SCC selection, improving the likelihood of isolating robust, high-yielding clones. The stable integration of the trastuzumab transgene with the Harbor-IN transposase and insulated transposon vectors provided integration at open chromatin and protection from gene silencing, further enhancing expression stability and productivity.

Finally, we enabled the identification of the expected molecular masses for the light and heavy chains after trastuzumab purification extracted from CleanCut GS clone, as well as the characterization of glycoform variants present on the heavy chain. As anticipated, glycan-dependent mass heterogeneity was observed for the heavy chain, whereas no glycan mass was detected on the light chain (Figure 9S). Together, the reduced LC–MS data confirmed the integrity, identity, and relative abundance of both antibody chains, providing a molecular-level characterization of trastuzumab.

## 5. Conclusion

Collectively, our findings highlight the value of Cas-CLOVER, CleanCut GS, and Harbor-IN as practical tools for industrial cell line development. Unlike Cas9, which is constrained in commercial CLD, Cas-CLOVER provides high-efficiency, high-specificity editing uniquely suited for GMP manufacturing platform creation. When combined with the Harbor-IN transposase, this system enables the rapid generation of stable, high-yielding CHO cell lines without downstream licensing burdens. Looking ahead, we continue to use Cas-CLOVER to expand the CleanCut GS platform by combining it with additional beneficial knockouts such as glycoengineering for enhanced antibody-dependent cell cytotoxicity (ADCC). We are also exploring the multiplexing potential of Cas-CLOVER to target problematic host cell proteins (HCPs) and pushing its boundaries on targeted integration capability. Importantly, our study employed a straightforward, non-optimized deep-well plate process using a commercially available medium. Thus, it is expected that GMP manufacturers applying optimized fed-batch or other advanced processes would achieve even higher titers and productivity with the easy to adopt CleanCut GS and Harbor-IN platform.

## Supporting information

Cintia Gomez Limia et al_Supplementary Material

## Acknowledgements

Figures were created with BioRender.com

## Conflict of Interest

All authors are employees of Demeetra, there are patents and patent applications on these technologies.

## Data availability

Study findings are supported by data available upon request from the corresponding author.

