## Supplementary material for "Sequential, Chromosome-Specific Glutamine Synthetase Double Knockout with Cas-CLOVER Establishes Enhanced CHO Platforms for Cell Line Development": Cintia Gomez Limia et al_Supplementary Material: Cintia Gomez Limia et al_Supplementary Material.docx

^1^ Demeetra AgBio, Lexington, Kentucky, 40505, USA.

**Supplementary Material for Review**

**Material and methods**

1.1. **Flow Cytometry analysis**

FACSymphony™ A3 (BD Bioscience) was used to evaluate the morphology and GFP expression of CHO cells. Cells (3 x 10^5^ cells) were harvested 72 h post transfection and resuspended in 70 uL of PBS. For viability analysis, cells were stained with 7AAD (Thermo Fisher Scientific, Cat# 00-6993-50) following the manufacturer’s instructions.

1.2. **PCR for genotyping validation**

PCR products were resolved on agarose gels, stained with ethidium bromide, and visualized to estimate band sizes. The amplicons were purified using magnetic beads Axygen® AxyPrep MAG PCR Clean-Up following manufacturer’s instructions (Corning, Cat# 16424001). The primers used for genotyping are listed in Table 2.

1.3. **qPCR for GS gene expression**

Total RNA was isolated from cells pellet using RNA extraction kit (Cat#T2110, NEB) according to the manufacturer’s instructions. Complementary DNA (cDNA) was synthesized from 500 ug total RNA using the SuperScript™ IV VILO™ cDNA Synthesis Kit VILO (Thermo Fisher Scientific, Cat#11766050) following the manufacturer’s protocol. Gene expression was assessed using TaqMan™ Gene Expression Assays (Thermo Fisher Scientific) with the following probe/assay IDs: GAPDH (Cg04424038_gH) and glutamine synthetase (GS) (Cg04424042_sH). Relative gene expression was calculated using the 2^-ΔΔCt method. Ct values for target genes were normalized to GAPDH (ΔCt) as reference gene, and fold changes were calculated relative to GS-expressing CHO cell. Experiments were performed with 2 independent replicates. Data is presented as mean ± SD.

1.4. **Trastuzumab purification**

Culture supernatants were clarified by centrifugation at 125 × g for 7 min to remove the cell and debris from the cell culture. The supernatant was then centrifuged at 2000g for 3 min and filtered through a 0.22-µm syringe filter. One milliliter of resin was washed thrice with 10 mL PBS and mixed with the filtered supernatant, followed by rocking incubation at room temperature for 30 min. The resin was pelleted, and the supernatant was labeled as flow-through and collected for further characterization. The resin was then washed six times with 10 mL PBS, with the first two washes saved as W1 and W2. While washing, six microcentrifuge tubes were prepared with 50 uL neutralizing buffer (1M Tris pH8.5). Elution was performed by adding 450 µL elution buffer to the resin, gently resuspend the pellet, and spun down at 2000g for 3 min. The supernatant was carefully removed and added to previously prepared tubes with 50 uL neutralization buffer without disturbing the resin pellet. It was then repeated 5 more times. All collected fractions were pulled together and analyzed by SDS-PAGE to assess protein recovery and purity.

1.5. **Reduced LC–MS Analysis of Trastuzumab**

Reduced LC–MS analysis was performed to identify and characterize the mass of light and heavy chains of trastuzumab. Purified antibody samples were prepared by diluting them in ammonium bicarbonate buffer. They were then treated with a reducing agent to reduce the intermolecular disulfide bonds, yielding fully separated light and heavy chains. The reduced samples were subsequently injected into LC-MS. The heavy chain and light chain were separated through a C4 column and analyzed by a high-resolution MS spectrum.

The heavy chain and light chain of Trastuzumab was separated by a liquid chromatography Li on a C4 reverse-phase column using a gradient of acetonitrile with 0.1% formic acid. Under these separation conditions, the light chain eluted earlier than the heavy chain due to differences in hydrophobicity. The eluate was introduced into a high-resolution mass spectrometer operating in positive electrospray ionization mode. Mass spectra were acquired across an extended m/z range suitable for intact protein detection. The resulting spectra were deconvoluted using standard intact-mass analysis software to generate zero-charge mass profiles.

**Tables**

**Table S1. Site candidates list for off-Target screening in GS5-SKO and GS-DKO clones after Cas-CLOVER editing.**

| **Target** | **gRNA** | **Site*^a^*** | **Location (Chr)** | **Start** | **End** |
| --- | --- | --- | --- | --- | --- |
| GS5 | gRNA-L | Glul-Left-Site-1 | NW_003614555.1 | 174626 | 175648 |
|  |  | Glul-Left-Site-2 | NW_003615010.1 | 262334 | 263356 |
|  |  | Glul-Left-Site-3 | NW_003613794.1 | 1282950 | 1283972 |
|  |  | Glul-Left-Site-4 | NW_003618130.1 | 28623 | 29645 |
|  |  | Glul-Left-Site-5 | NW_003614676.1 | 96571 | 97593 |
|  |  | Glul-Left-Site-6 | NW_003614345.1 | 253216 | 254238 |
|  |  | Glul-Left-Site-7 | NW_003616279.1 | 72724 | 73746 |
|  |  | Glul-Left-Site-8 | NW_003613710.1 | 295449 | 296471 |
|  |  | Glul-Left-Site-9 | NW_003614356.1 | 508948 | 509970 |
|  |  | Glul-Left-Site-10 | NW_003613902.1 | 1228501 | 1229523 |
|  | gRNA-R | Glul-Right-Site-1 | NW_003616279.1 | 72764 | 73786 |
|  |  | Glul-Right-Site-2 | NW_003613948.1 | 693287 | 694309 |
|  |  | Glul-Right-Site-3 | NW_003615161.1 | 259672 | 260694 |
|  |  | Glul-Right-Site-4 | NW_003614830.1 | 177895 | 178917 |
|  |  | Glul-Right-Site-5 | NW_003614076.1 | 598138 | 599160 |
|  |  | Glul-Right-Site-6 | NW_003614242.1 | 93742 | 94764 |
|  |  | Glul-Right-Site-7 | NW_003614364.1 | 628722 | 629744 |
|  |  | Glul-Right-Site-8 | NW_003614363.1 | 514570 | 515592 |
|  |  | Glul-Right-Site-9 | NW_003613852.1 | 1118917 | 1119939 |
|  |  | Glul-Right-Site-10 | NW_003613596.1 | 1908910 | 1909932 |
| GS1 | gRNA-L | Pseudo-Left-Site-1 | NW_003613921.1 | 1433106 | 1434128 |
|  |  | Pseudo-Left-Site-2 | NW_003616444.1 | 17279 | 18301 |
|  |  | Pseudo-Left-Site-3 | NW_003613736.1 | 2123702 | 2124724 |
|  |  | Pseudo-Left-Site-4 | NW_003613835.1 | 1782774 | 1783796 |
|  |  | Pseudo-Left-Site-5 | NW_003613880.1 | 311040 | 312062 |
|  |  | Pseudo-Left-Site-6 | NW_003614008.1 | 857404 | 858426 |
|  |  | Pseudo-Left-Site-7 | NW_003617724.1 | 35188 | 36210 |
|  |  | Pseudo-Left-Site-8 | NW_003613629.1 | 3691537 | 3692559 |
|  |  | Pseudo-Left-Site-9 | NW_003615422.1 | 131913 | 132935 |
|  |  | Pseudo-Left-Site-10 | NW_003614094.1 | 189417 | 190439 |
|  | gRNA-R | Pseudo-Right-Site-1 | NW_003613921.1 | 1433606 | 1434628 |
|  |  | Pseudo-Right-Site-2 | NW_003616279.1 | 72474 | 73496 |
|  |  | Pseudo-Right-Site-3 | NW_003614196.1 | 651894 | 652916 |
|  |  | Pseudo-Right-Site-4 | NW_003613726.1 | 52019 | 53041 |
|  |  | Pseudo-Right-Site-5 | NW_003614646.1 | 156273 | 157295 |
|  |  | Pseudo-Right-Site-6 | NW_003614289.1 | 204289 | 205311 |
|  |  | Pseudo-Right-Site-7 | NW_003613836.1 | 972399 | 973421 |
|  |  | Pseudo-Right-Site-8 | NW_003613676.1 | 1748962 | 1749984 |
|  |  | Pseudo-Right-Site-9 | NW_003614474.1 | 70291 | 71313 |
|  |  | Pseudo-Right-Site-10 | NW_003613756.1 | 1593253 | 1594275 |

*^a^*The sites listed correspond with the 23bp CRISPOR off-Target sequences with 500bp pads upstream (Start) and downstream (End).

**Table S2. Variants were identified in CHO-K1 WT, 7G2 and CleanCut GS.**

| **Target** | **gRNA** | **Off-Target Site^a^** | **Location (Chr)** | **Position^b^** | **Allelic Frequency^b^** | **Distance (bp) from site^b^** | **Type**  **of variant^c^** |
| --- | --- | --- | --- | --- | --- | --- | --- |
| GS5 | gRNA-L | Glul-Left-Site-8 | NW_003613710.1 | 295671 | 0.5 | 289 | Ins |
|  |  | Glul-Left-Site-10 | NW_003613902.1 | 1228441 | 1 | 571 | mnp |
|  | gRNA-R | Glul-Right-Site-4 | NW_003614830.1 | 178240 | 0.5 | 166 | SNP |
|  |  |  |  | 178449 | 0.5 | 43 | SNP |
|  |  | Glul-Right-Site-5 | NW_003614076.1 | 598802 | 0.5 | 153 | SNP |
|  |  | Glul-Right-Site-6 | NW_003614242.1 | 93932 | 0.5 | 321 | Ins |
|  |  | Glul-Right-Site-7 | NW_003614364.1 | 629648 | 1 | 415 | mnp |
|  |  |  |  | 629666 | 1 | 433 | SNP |
|  |  |  |  | 629799 | 1 | 566 | SNP |
|  |  |  |  | 629844 | 1 | 611 | SNP |
| GS1 | gRNA-L | Pseudo-Left-Site-2 | NW_003616444.1 | 17525 | 0.5 | 265 | SNP |
|  |  |  |  | 17694 | 0.5 | 96 | SNP |
|  |  |  |  | 17861 | 0.5 | 71 | SNP |
|  |  |  |  | 17868 | 0.5 | 78 | SNP |
|  |  |  |  | 18213 | 0.5 | 423 | SNP |
|  |  | Pseudo-Left-Site-3 | NW_003613736.1 | 2123811 | 0.5 | 402 | SNP |
|  |  |  |  | 1783731 | 1 | 446 | SNP |
|  |  | Pseudo-Left-Site-4 | NW_003613835.1 | 1783750 | 1 | 465 | SNP |
|  |  |  |  | 1783756 | 1 | 471 | SNP |
|  |  | Pseudo-Left-Site-7 | NW_003617724.1 | 35382 | 0.5 | 317 | Del |
|  |  |  |  | 35828 | 0.5 | 129 | Del |
|  |  |  |  | 35865 | 1 | 166 | mnp |
|  | gRNA-R | Pseudo-Right-Site-3 | NW_003614196.1 | 652263 | 1 | 142 | SNP |
|  |  |  |  | 652829 | 1 | 424 | mnp |
|  |  | Pseudo-Right-Site-6 | NW_003614289.1 | 204304 | 0.5 | 496 | SNP |
|  |  |  |  | 204420 | 0.5 | 380 | SNP |
|  |  |  |  | 204663 | 0.5 | 137 | SNP |
|  |  |  |  | 204880 | 0.5 | 80 | Del |
|  |  |  |  | 204973 | 0.5 | 173 | SNP |
|  |  |  |  | 205354 | 0.5 | 554 | Ins |
|  |  | Pseudo-Right-Site-10 | NW_003613756.1 | 1593471 | 0.5 | 293 | SNP |

^a^The three samples were compared with the Reference sequence for GS5 and GS1 using the off-target sites.

^b^The position, allelic Frequency, distance and type of variant are described per each off-target site that showed difference with the Reference sequence.

^c^Type of variants is described as Ins: Insertion, del: deletion, SNP: Single-nucleotide polymorphism, MNP: multiple-nucleotide polymorphism.

**Table S3. On-Target Site analysis in the GS5 and GS1 loci comparing CHO-K1 WT, 7G2 and CleanCut GS clones.**

| **Target** | **CHO-K1 WT presented variants** | **7G2 presented variants** | **CleanCut GS present variants** | **Location^a^** | **Position** | **Allelic Frequency** | **Type of Variant^a^** | **Distance from Site** |
| --- | --- | --- | --- | --- | --- | --- | --- | --- |
| GS5 | N | Y | Y | NW_003613921.1 | 1434554 | 0.5,0.5 | del, del | 14 |
| GS1 | Y | Y | N | NW_003614063.1 | 177086 | 0.5 | SNP | 542 |
|  | N | N | Y | NW_003614063.1 | 177631 | 1 | SNP | 3 |
|  | Y | Y | Y | NW_003614063.1 | 177715 | 0.5 | SNP | 87 |
|  | Y | Y | Y | NW_003614063.1 | 177816 | 0.5 | SNP | 188 |

^a^The locations, allelic frequency and type of variant were summarized for the cell line that is carrying the modification compared with the genome reference used for the analysis. Variants present is indicated by Yes (Y) or Not (N).

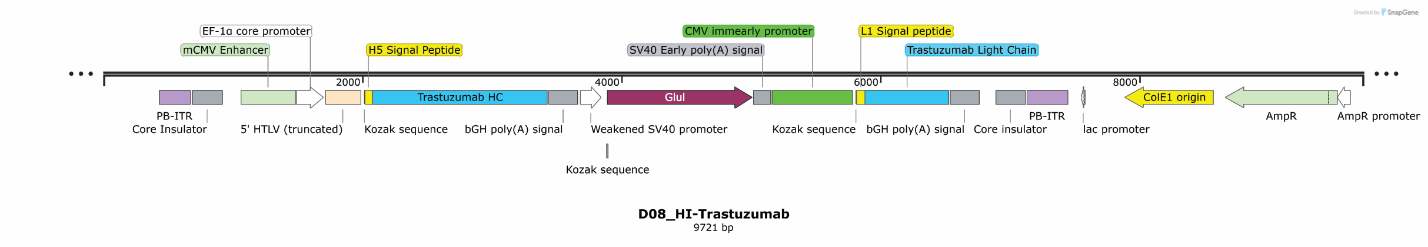
**Figures**

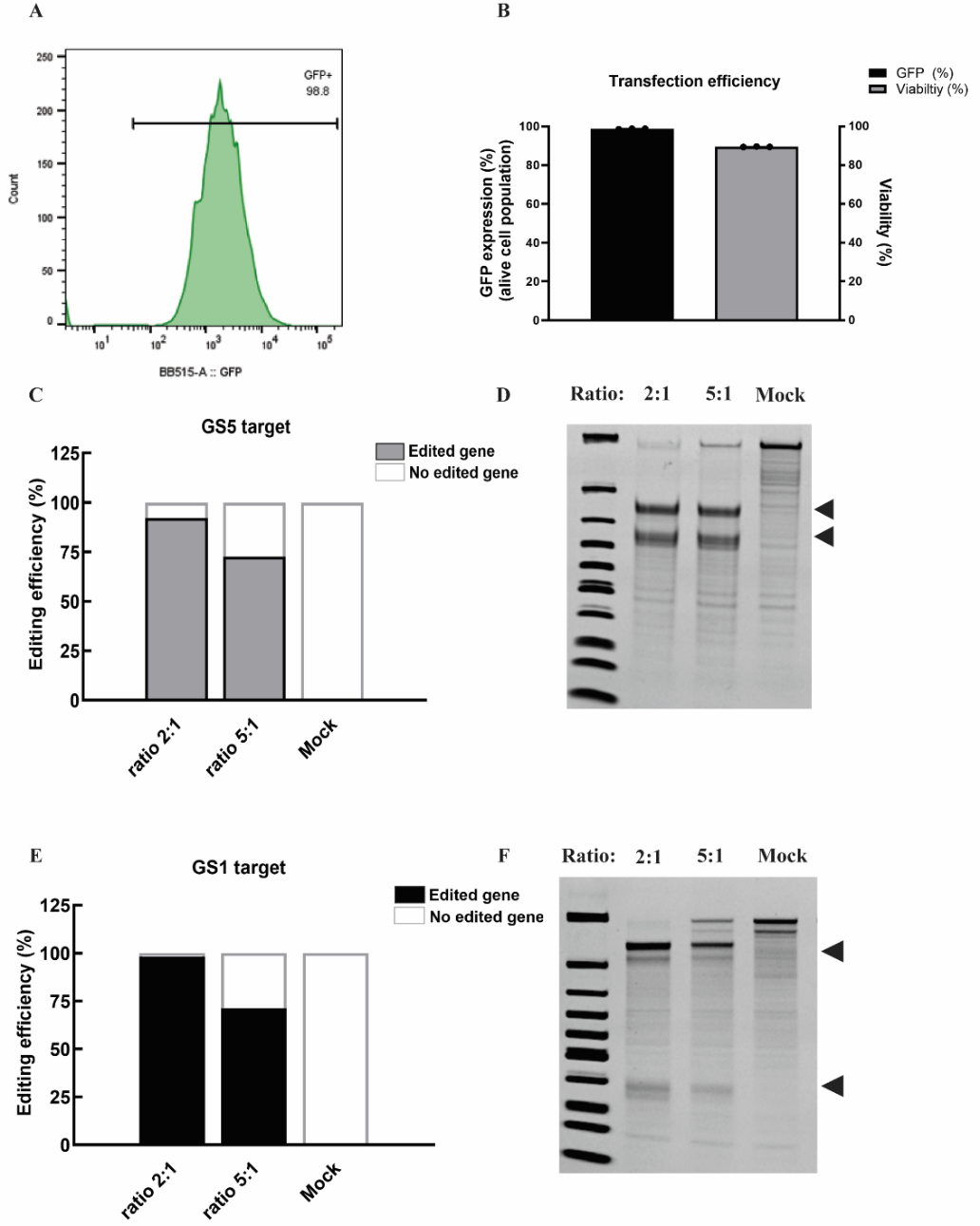
**Figure S1. Construct encoding trastuzumab in the Harbor-IN transposon.** Trastuzumab heavy chain is expressed using a murine CMV (MCMV) major immediate-early enhancer, an Ef1α core promoter, and a truncated 5′ HTLV-1 LTR. The trastuzumab light chain is driven by the human CMV (HCMV) immediate-early enhancer/promoter. GS selection marker is regulated by a weakened SV40 promoter.

**Figure S2. Transfection efficiency and gene editing validation for GS5 and GS1 targets.**
**(A)** Flow cytometry histogram showing GFP expression. Experiments were performed in triplicates, and error bars represent mean ± SD. **(B)** Quantification of transfection efficiency measured as GFP expression (black bars, left y-axis) and cell viability (gray bars, right y-axis). **(C, E)** Different Cas-CLOVER:gRNA ratios (2:1 and 5:1) were tested, with mock-transfected cells serving as negative controls for GS5 and GS1 respectively. **(D,F)** Representative images of PCR amplicons from GS5 and GS1 by T7 assay targets following Cas-CLOVER editing. Black arrowheads indicate edited fragments. No editing was observed in control conditions (Cas-CLOVER (CC) – no gRNA). For GS5 target: the edited fragments are 413 bp and 339 bp from the unedited 752 bp band. For GS1 target: the edited fragments are in 618 bp and 152 bp from the unedited 770 bp band.

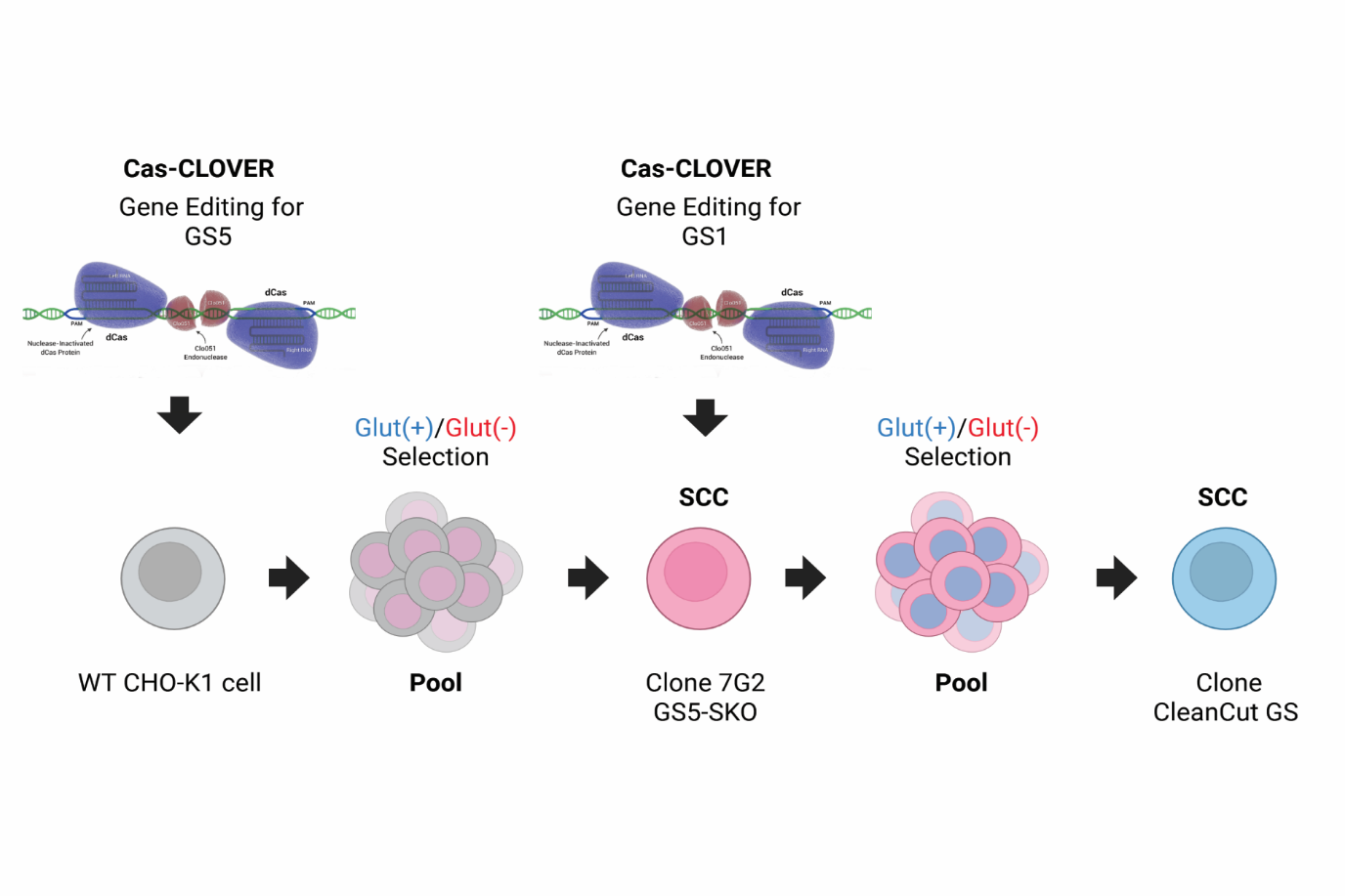

**Figure S3. Generating a CHO GS5-SKO or GS-DKO cell lines.** The schematic representation shows the steps to generate the GS KO cell lines via Cas-CLOVER. A parental WT CHO-K1 cell line was used to generate the GS5-SKO pool cells targeting the *GS5* locus. The top clone 7G2 was edited to target GS1 and generate the GS-DKO pool cells. From this GS-DKO pool a top clone F8 was isolated with a confirmed GS-DKO genotype. To evaluate glutamine sensitivity, cell pools and later isolated clones were grown in CHO medium with L-glutamine (+) (highlighted in blue) or without (–) L-glutamine (highlighted in red). Glut: L-glutamine, SSC: Single Cell Clone.

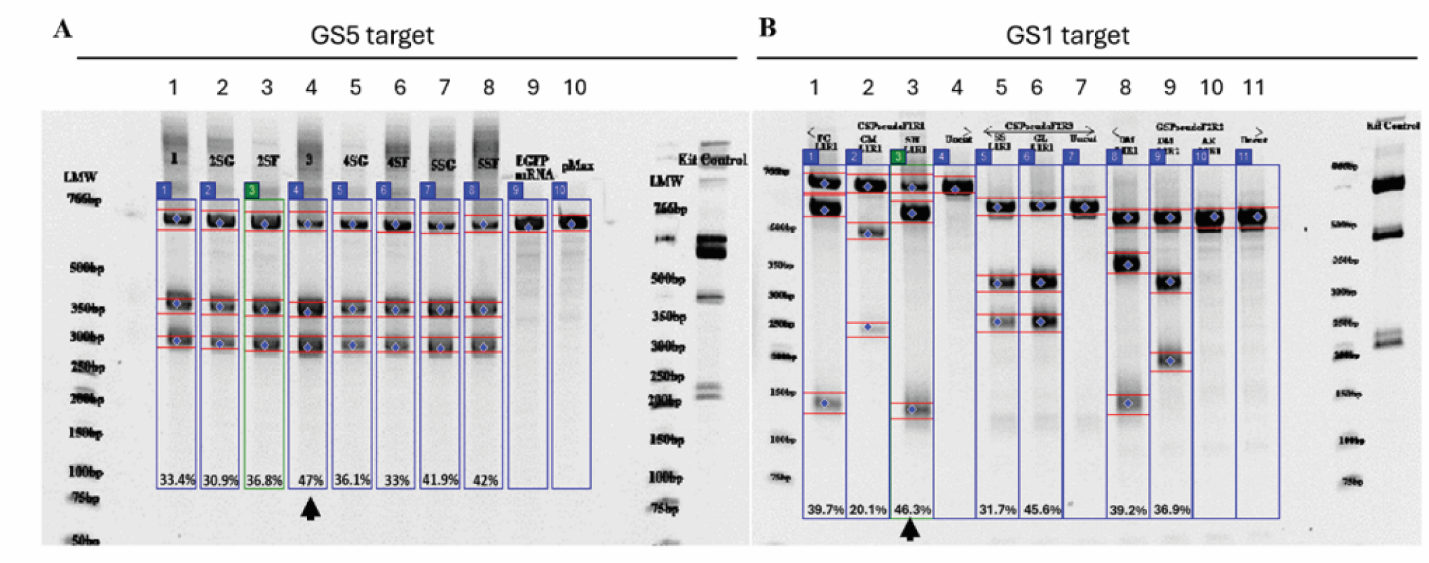
**Figure S4. Cas-CLOVER–mediated editing of *GS* targets in pooled CHO cell analyzed by T7 Endonuclease I assay. (A)** 4–20% TBE polyacrylamide gel showing T7 EnGen® (T7 Endonuclease I) assay results from CHO cell pools transfected with Cas-CLOVER mRNA and sgRNAs targeting GS5 loci. Lane 4 is corresponding with the CHO-K1 WT cells nucleofected to target GS5 loci. Low Molecular Weigth is the DNA ladder, and control corresponds with lanes 9 and 10, EGFP mRNA, pMax are included for reference. **(B)** Corresponding assay of pools targeting GS1 loci. Each numbered lane represents an individual guide pair or condition, with the measured indel percentage indicated below. Black arrows mark the guide pair/condition selected for further clonal isolation and characterization. Lane 3 is corresponding with the CHO SKO cells nucleofected to target GS1 loci.

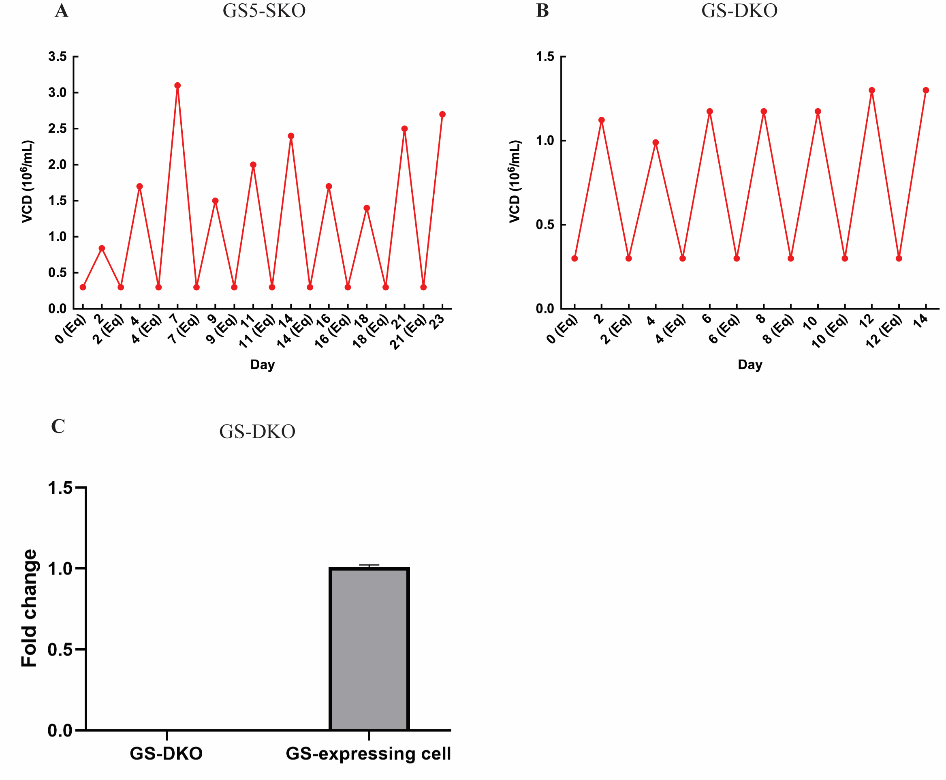

**Figure S5.** **Viable cell density (VCD) during serial passages GS5-SKO and GS-DKO GS expression analysis. (A, B)** Growth profile of 7G2 or CleanCut GS clone, receptively in repeated batch culture. **C.** mRNA level analysis for GS expression in CleanCut GS clone.

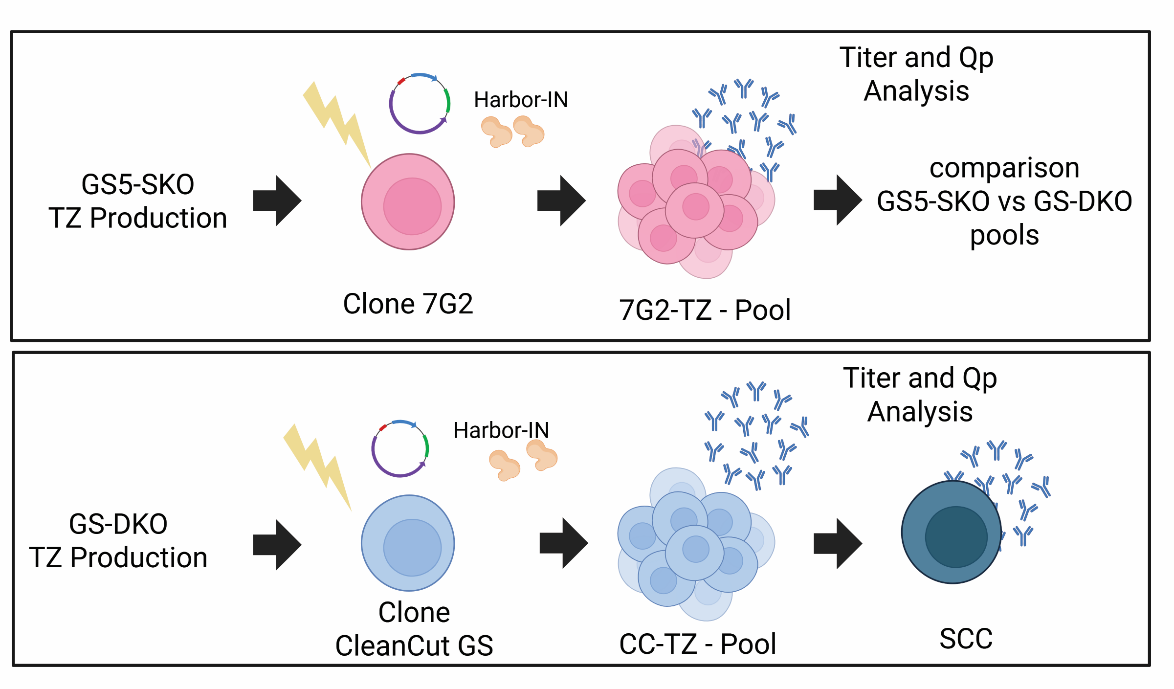

**Figure S6.** **Generation of GS5-SKO and GS-DKO CHO cell lines and Trastuzumab-expressing pool or clones using Cas-CLOVER and Harbor-IN transposase technology.** Clone 7G2 was selected to transfect with Harbor-IN transposase and transposon encoding TZ (Top). Clone CleanCut GS was selected to transfect with Harbor-IN transposase and a transposon encoding TZ. After SCC, clones were isolated from this pool (Bottom). Pools and isolated clones were evaluated for Trastuzumab titers and specific productivity (Qp) values. CC-TZ: CleanCut GS clone.

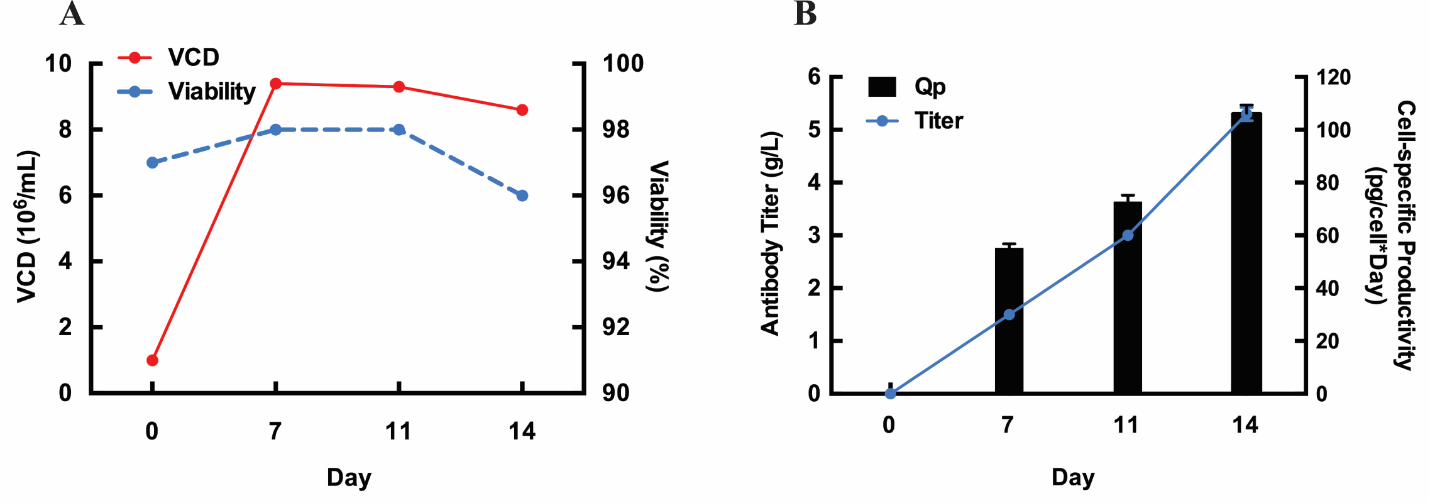

**Figure S7. Characterization of GS-DKO CHO cell expressing Trastuzumab. (A)** Viable cell density (VCD) and viability of the 3h31-TZ clone for a 14-day culture. **(B)** Titer and specific productivity (Qp) of 3H31-TZ for 14 days culture. Experiments were performed in triplicates, and error bars represent mean ± SD.

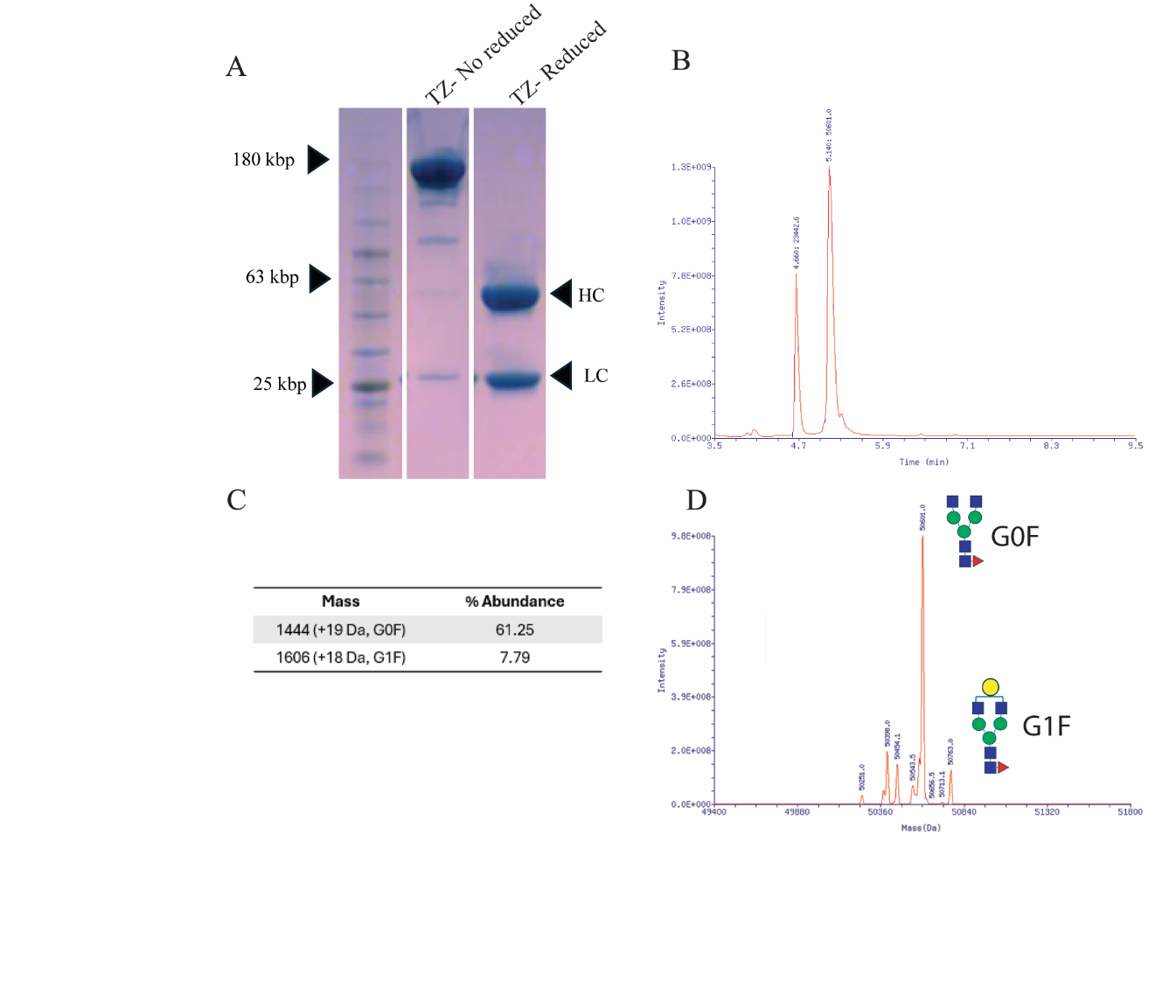

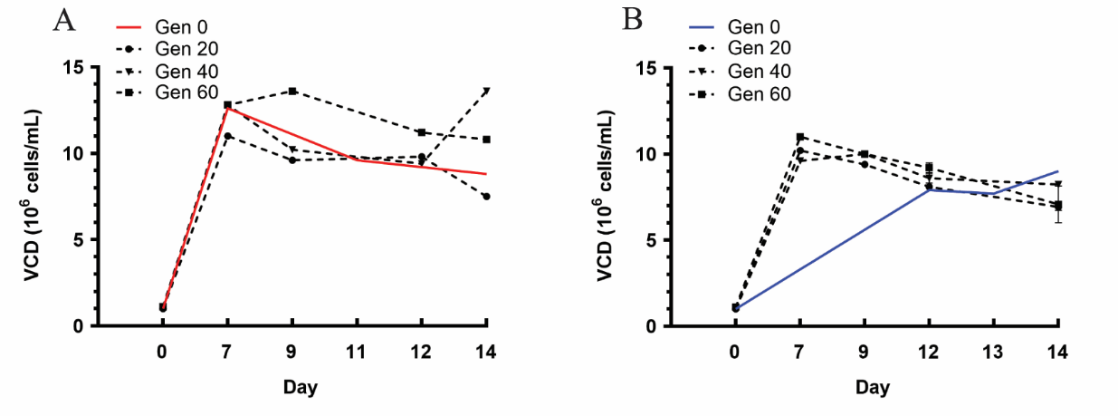
**Figure S8. Stability of CleanCut GS-TZ pool and 5G2-TZ clone over 60 generations. (A, B)** VCD measured in CleanCut GS-TZ pool and 5G2-TZ clone over 20, 40 and 60 generations and compared with Generation 0 for 14 days. Experiments were performed in triplicates, and error bars represent mean ± SD.

**Figure S9. Analysis of protein-level and glycan profile of trastuzumab produced by the GS-DKO-TZ pool. (A)** SDS–PAGE analysis of purified trastuzumab under reducing conditions confirms the presence of both light chain (LC, ~25 kDa) and heavy chain (HC,~50 kDa), indicating correct antibody assembly and purity following Protein A–based purification. Gel was stained with Comises Blue. Ladder is 1kb band. **(B)** Reduced reverse-phase LC chromatogram of trastuzumab showing clear separation of the LC and HC. The LC elutes at 4.66 min and the HC at 5.14 min. Integrated peak areas demonstrate an approximate 1:2 LC: HC ratio, consistent with the expected molecular weight difference and stoichiometry of an IgG molecule. Deconvoluted reduced LC–MS spectrum of the LC showing a single dominant mass corresponding to the unglycosylated LC, with no detectable glycan-associated mass heterogeneity. **(C)** Deconvoluted reduced LC–MS spectrum of the HC highlighting glycan-dependent mass variants. **(D)** The predominant glycoforms correspond to G0F (61.25%) and G1F (7.7%), confirming Fc glycosylation typical of trastuzumab.
